# Disease-linked mutations dysregulate neuronal condensate physical properties, composition, and RNA translation

**DOI:** 10.1101/2024.11.01.621623

**Authors:** Leshani Ahangama Liyanage, Fraser McCready, Steve Chung, Jason Arsenault, Wei Wei, Xusheng Lin, Lu-Yang Wang, James Ellis, Jonathon A. Ditlev

**Author notes:** Corresponding Author (J.A.D).

## Abstract

Local RNA translation is essential for development. In neurons, deficient local translation linked with mutations in scaffold proteins results in dysregulated dendrite and dendritic spine growth. However, mechanisms by which these proteins control translation and how disease-linked mutations induce aberrant translation were unclear. We use biochemical reconstitution and neuronal assays to show that mutations to the neuronal condensate scaffold shank2 cause physical hardening and altered composition of condensates; a key RNA translation-modulator FMRP is excluded from mutant condensates. Functionally, shank2 condensates repress translation while condensates composed of shank2 with intrinsically disordered region-localized missense mutations promote translation. These results demonstrate that disease-linked dysregulation of condensate physical properties and composition is an underlying mechanism of aberrant RNA translation often observed in disease.

**One Sentence Summary:** Disease-linked missense mutations in the postsynaptic density scaffold protein shank2 dysregulate phase separated biomolecular condensate physical properties and composition resulting in aberrant RNA translation.

## Introduction

Biomolecular condensates are ubiquitous structures that functionally organize proteins and nucleic acids without encapsulating membranes (*1*). Many condensates form through the process of biological phase separation and can spatiotemporally control specific cellular functions (*2–4*). In neurons, spatiotemporal control of cellular functions, such as RNA translation, is especially important because of their highly polarized cell structures that includes long projections, i.e., dendrites and axons. To accomplish local functions in polarized regions, neurons package RNA into transport granules where their local translation can be modulated by other condensates, including those formed by postsynaptic density (PSD) scaffold proteins (*5*). Phase separation of PSD scaffold proteins has been previously linked to control of the actin cytoskeleton and the spatial organization of proteins, including kinase and membrane receptor enrichment and sorting of excitatory and inhibitory synapse proteins (*6*, *7*).

The PSD scaffolding protein shank2 modulates neuron dendrite growth and connectivity (*8*, *9*). Disease-linked missense mutations in shank2 result in an altered FMRP-modulated transcriptome (*8*) and protein expression (*10*), suggesting functional interactions between shank2 and FMRP. This is especially important in neurons because the expression of many neuron-specific proteins responsible for dendrite growth and connectivity are under the control of Fragile X Mental Retardation Protein (FMRP) (*11*). Shank2 deficiencies ultimately result in a hyperactive network of neurons that deviate from isogenic wild type (WT) control networks (*8*, *9*). A recent study identified disease-linked mutations in shank2 that are predicted to alter its phase separation, suggesting a link between aberrant phase separation and mechanisms of disease (*12*). However, neither the mechanism by which shank2 controls local RNA translation nor the cause of aberrant FMRP-dependent translation downstream of shank2 mutations are known.

### Shank2 condensates are not dynamic and repress mRNA translation

The domains of shank-family proteins are sufficient for their phase separation, however the role of their often-mutated ∼1000 residue intrinsically disordered region (IDR) in phase separation has not been evaluated (*6*). Mutations in shank2 domains have been linked with deficient shank2 nanocluster formation (*13*); however the role of the shank family IDR in modulating phase separation was elusive. To define the contribution of the shank2 IDR to its phase separation, we purified full-length shank2a (hereafter shank2) and found that it coalesces into micron-sized biomolecular condensates with its binding partner homer1 at nanomolar concentrations (Fig. 1A; fig. S1), ∼50-fold lower than fragments of shank family proteins consisting of only folded domains and binding motifs likely due to IDR-IDR interactions between shank2 proteins (*6*). Consistent with phase separation, condensate formation depends on the concentrations of shank2 and homer1 (Fig. 1B). We probed the dynamics of shank2 within droplets using fluorescence recovery after photobleaching (FRAP) and found that within ∼1 min of formation, shank2 dynamics are arrested (Fig. 1C). These results suggest that following spherical condensate formation typical of liquid-like phase separation, a rapid physical transition into a gel-like state occurs (*14*, *15*). Previous work demonstrates that phosphorylation of phase-separated IDRs can prevent physical solid-to-liquid phase transitions of condensed proteins (*16*). To determine whether shank2 phosphorylation alters protein dynamics, we phosphorylated shank2 (pshank2) using mitogen activated protein kinase, confirmed phosphorylation using mass spectrometry (fig. S2), and repeated the above experiments. Phosphorylation of shank2 neither promoted changes in its propensity to phase separate with homer1 (Fig. 1D; fig. S3) nor altered its dynamics within condensates (Fig. 1E). In a neuronal cell line, shank2 mutations have been linked with dysregulated protein expression (*10*). To test whether the shank2 condensates can affect RNA translation, we performed *in vitro* RNA translation assays containing shank2 condensates (*17*). We found that the presence of shank2 or pshank2 condensates resulted in a ∼5 order of magnitude reduction in RNA translation compared to a control with no shank2 condensates (Fig. 1F). Importantly, shank2 condensate formation in lysate (fig. S4) was similar to what was observed in solution (Fig. 1A, D), allowing us to directly link shank2 condensates with dampened RNA translation *in vitro*.

**Fig. 1.**
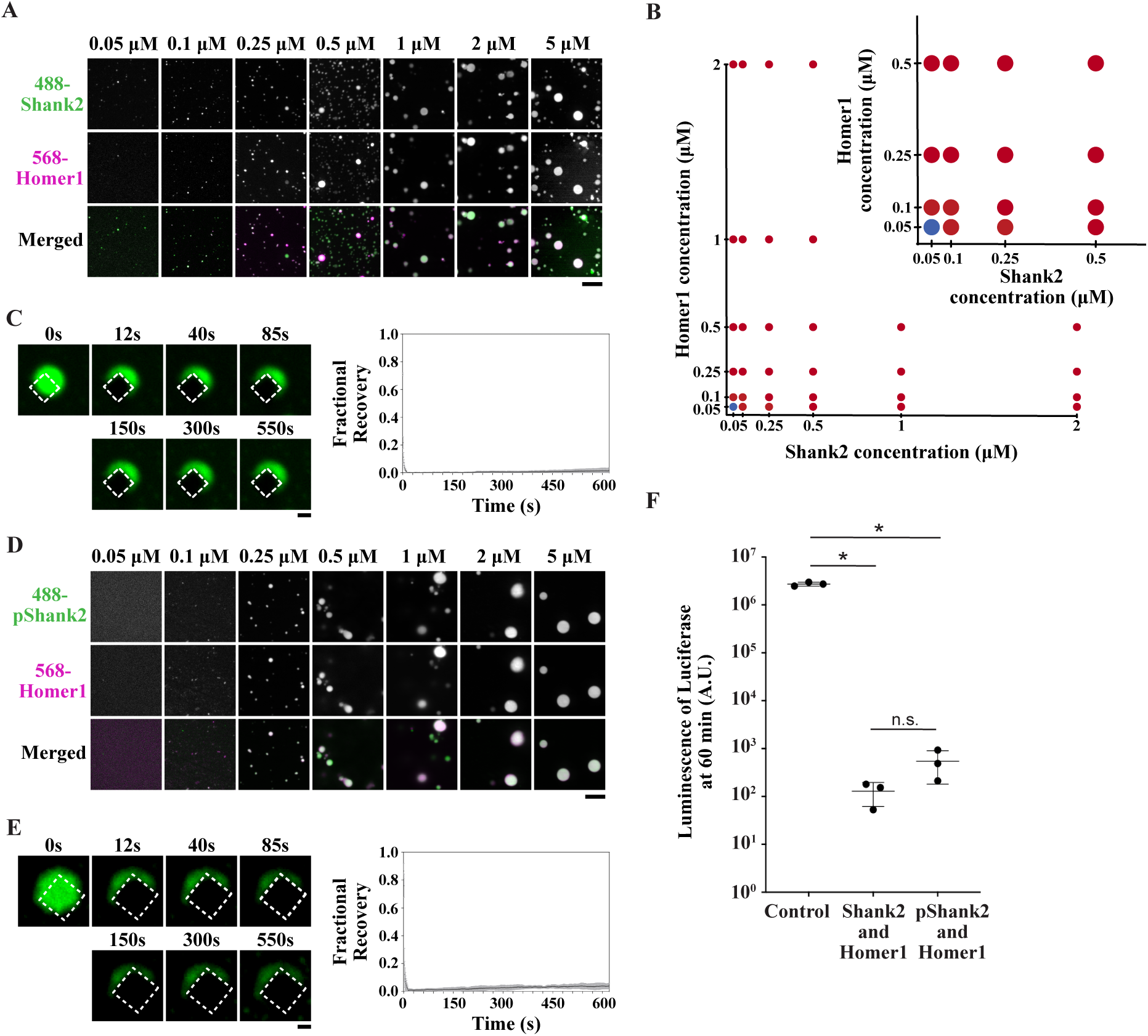
Phase separated shank2 and homer1 condensates significantly dampen RNA translation *in vitro*. **(A)** Confocal fluorescence microscopy imaging of AlexaFluor (AF) 488-shank2 (green) and AF568-homer1 (magenta) condensation from 0.05 μM to 5 μM. Below 0.05 μM, visible condensates did not form. Scale bar = 10 μm. N = 3 experiments with 5 fields of view each analyzed. **(B)** Phase diagram of shank2 and homer1 as concentrations of each increased from 0.05 μM to 5 μM. Blue dots indicate no observable phase separation; red dots indicate phase separation. N = 3 experiments with 5 fields of view each analyzed. **(C)** Fluorescence recovery after photobleaching (FRAP) of AF488-shank2 (green) and homer1 condensate. Time 0 s was captured before photobleaching, 12 s was captured immediately after the photobleaching pulse. Dashed white box illustrates region of photobleach. Scale bar = 1 μm. Plot on bottom shows the time course of recovered for AF488-shank2 in condensates formed with shank2 and homer1. Shown are the mean (black line) ± s.d. (gray region above and below black line). N = 3. **(D)** Confocal fluorescence microscopy imaging of phosphorylated AF488-shank2 (green; pshank2) and AF568-homer1 (magenta) condensation from 0.05 μM to 5 μM. Below 0.05 μM, visible condensates did not form. Scale bar = 10 μm. See fig. S3 for full phase diagram across pshank2 and homer1 concentrations from 0.05 μM to 5 μM. N = 3 experiments with 5 fields of view each analyzed. **(E)** Fluorescence recovery after photobleaching (FRAP) of AF488-pshank2 (green) and homer1 condensate. Time 0 s was captured before photobleaching, 12 s was captured immediately after the photobleaching pulse. Dashed white box illustrates region of photobleach. Scale bar = 1 μm. Plot on bottom shows the time course of recovered for AF488-shank2 in condensates formed with shank2 and homer1. Shown are the mean (black line) ± s.d. (gray region above and below black line). N = 3. **(F)** *In vitro* translation of luciferase mRNA in rabbit reticulocyte lysate after 60 minutes. Control: lysate + luciferase mRNA. Shank2 and Homer1: lysate + shank2 and homer1 condensates + luciferase mRNA. pShank2 and Homer1: lysate + pshank2 and homer1 condensates + luciferase mRNA. Shown are mean ± s.d. N = 3. * = p < 0.05, t-test.

### FMRP is enriched in shank2 condensates

Previous proteomics work revealed that FMRP localizes to the PSD upon neuronal stimulation and the initiation of local mRNA translation (*18*). We therefore asked whether FMRP localizes to shank2/homer1 condensates. When we added the disordered C-terminal IDR tail of FMRP, we observed salt-dependent FMRP enrichment of FMRP in shank2/homer1 condensates (Fig. 2A). When solutions contained 150 mM NaCl, FMRP was not enriched in shank2/homer1 condensates; however, when solutions contained 50 mM NaCl, FMRP was highly enriched in shank2 / homer1 condensates. These data suggest that FMRP localizes to shank2 / homer1 condensates via electrostatic interactions. Phosphorylation of phase separating proteins increases their overall negative charge, which can reduce or increase protein enrichment in condensates by increasing electrostatic repulsion or attraction, respectively (*2*). We therefore tested the effect of phosphorylation on FMRP enrichment. Neither the phosphorylation of shank2 nor FMRP (pFMRP) resulted in significant changes to FMRP enrichment in condensates (Fig. 2B, C).

**Fig. 2.**
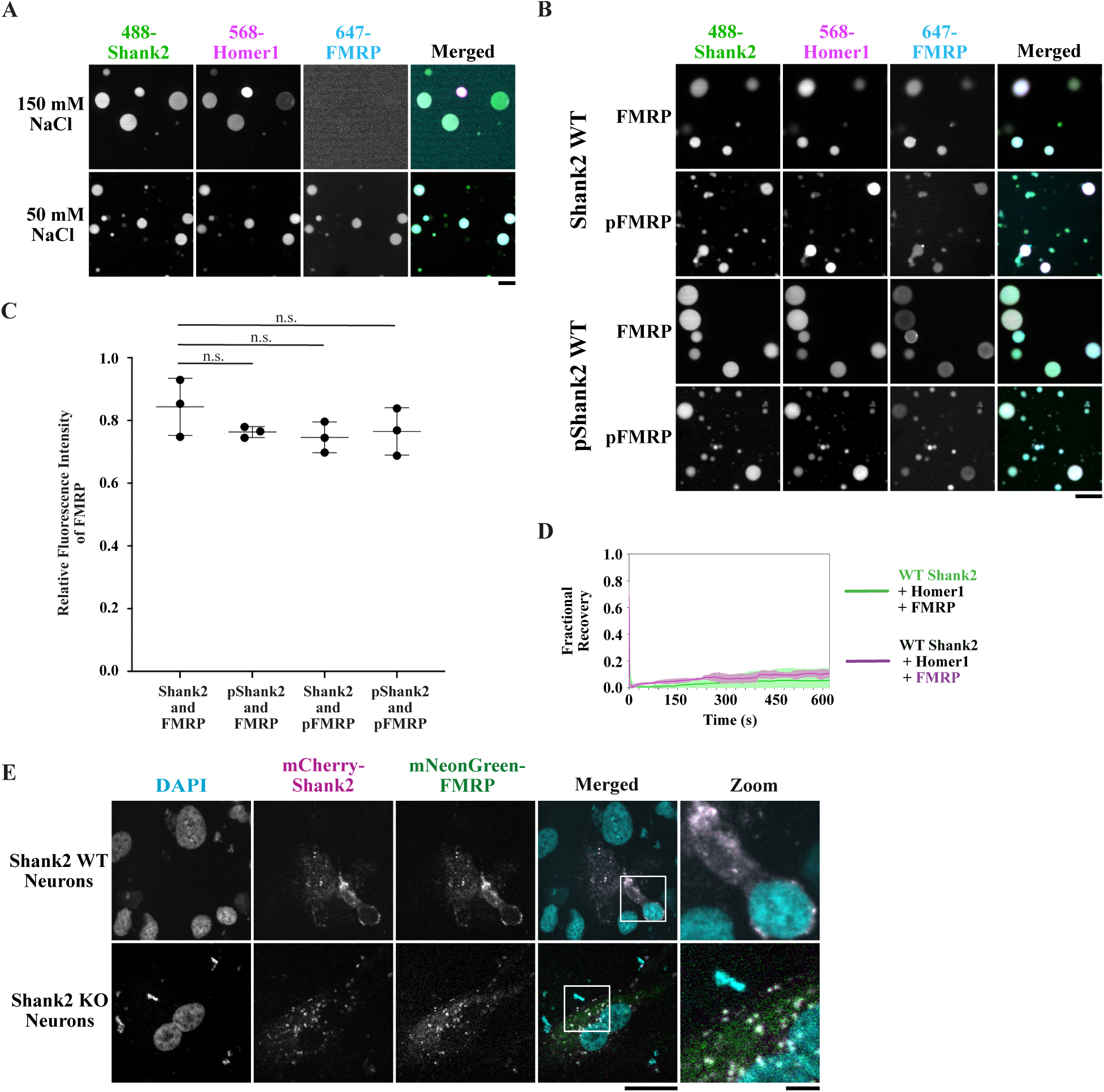
Shank2 condensates are enriched by FMRP. **(A)** Confocal fluorescence microscopy images of AlexaFluor (AF) 488-shank2 (green) and AF568-homer1 (magenta) condensates with AF647-FMRP (cyan). (top) In 150 mM NaCl buffer FMRP is not enriched in shank2 / homer1 condensates. (bottom) In 50 mM NaCl buffer FMRP is enriched in shank2 / homer1 condensates. Scale bar = 10 μm. N = 3 experiments with 5 fields of view each analyzed. **(B)** AF647-FMRP (cyan) or AF647-phospho-FMRP (pFMRP, cyan) are enriched in AF488-shank2 (green) / AF568-homer1 (magenta) condensates or AF488-phospho-shank2 (pshank2, green) / AF468-homer1 (magenta) condensates. Scale bar = 20 μm. N = 3 experiments with 5 fields of view each analyzed. **(C)** Quantification of images shown in **(B)**. Intensity of AF647-(p)FMRP was divided by intensity of AF488-(p)shank2 to normalize data across conditions. n.s. = not significant, t-test. **(D)** Fluorescence recovery after photobeaching curves of shank2 (green) and FMRP (magenta) shows the time course of recovery for shank2 and FMRP in condensates formed with shank2, homer1, and FMRP. Shown are the mean (green, magenta lines) ± s.d. (light green, light magenta region above and below line). N = 3. **(E)** Confocal fluorescence microscopy images of 4-week-old SMAD+ NPC-differentiated inactive shank2 wild-type (WT) or shank2 knock out (KO) human neurons transfected with mCherry-shank2 (magenta) and mNeonGreen-FMRP (green). Zoomed images are of white box regions of merged images. Scale Bar under merged images = 20 μm. Scale bar under zoomed images = 5 μm.

Additionally, enrichment of FMRP in shank2 or pshank2 condensates altered neither shank2 nor FMRP dynamics (Fig. 2D; fig. S5, S6). Taken together, these results indicate that FMRP enrichment in shank2 condensates depends on electrostatic interactions but is not significantly altered by phosphorylation. We then asked whether our reduced *in vitro* experimental system was predictive of a complex neuronal environment. We used isogenic WT and *SHANK2* homozygous knockout (KO) induced Pluripotent Stem Cells (iPSCs) to generate human cortical neurons at 4 weeks of differentiation (*8*, *9*) for co-transfection with mCherry-shank2 and full- length mNeonGreen-FMRP. We observed the formation of shank2-enriched condensates that colocalized with mNeonGreen-FMRP (Fig. 2E) in human neurons regardless of their endogenous shank2 levels, similar to our *in vitro* results.

### Disease-linked mutations in shank2 prevent FMRP enrichment

Because FMRP is well enriched in shank2-containing condensates *in vitro* and in iPSC- derived human neurons, we asked whether disease-linked mutations alter FMRP localization in these condensates. We selected missense mutations in the IDR of shank2 (R958S, G1170R, R1427W, A1731S) that i) result in change in charge (i.e., RtoS), ii) result in a new phosphorylatable residue (i.e., AtoS), and ii) have been clinically linked with autism spectrum disorders and schizophrenia (*19*, *20*). Using purified, mutated shank2, homer1, and the C- terminal tail of FMRP, we evaluated FMRP localization to condensates without and with phosphorylation. Strikingly, we observed that in experiments using R958S and A1731S pshank2, FMRP enrichment is significantly diminished compared to WT shank2 (Fig. 3A, B), while FMRP enrichment is not affected by RtoW or GtoR mutations (fig. S7). FMRP enrichment was diminished in R958S and A1731S pshank2 condensates across a 0.5 μM to 20 μM FMRP concentration range (Fig. 3C; fig. S8). Surprisingly, the saturation concentrations of R958S and A1731S pshank2 are similar to those of WT pshank2 (fig. S9). Moreover, neither R958S nor A1731S significantly altered shank2 phase separation nor shank2 dynamics in condensates (fig. S10, S11), although there was a slight increase in A1731S pshank2 fluorescence recovery at the edges of the condensates (fig. S11). The dynamics of FMRP in shank2 condensates is higher in shank2 A1731S compared to WT shank2 and R958S shank2 (Fig. 3D), although FMRP never approaches full fluorescence recovery. In our *in vitro* assays, we noted that FMRP was unable to penetrate into many of the mutant pshank2 condensates (Fig. 3A) even though the dynamics of shank2 molecules within condensates is not changed after ∼1 minute based on FRAP analysis.

**Fig. 3.**
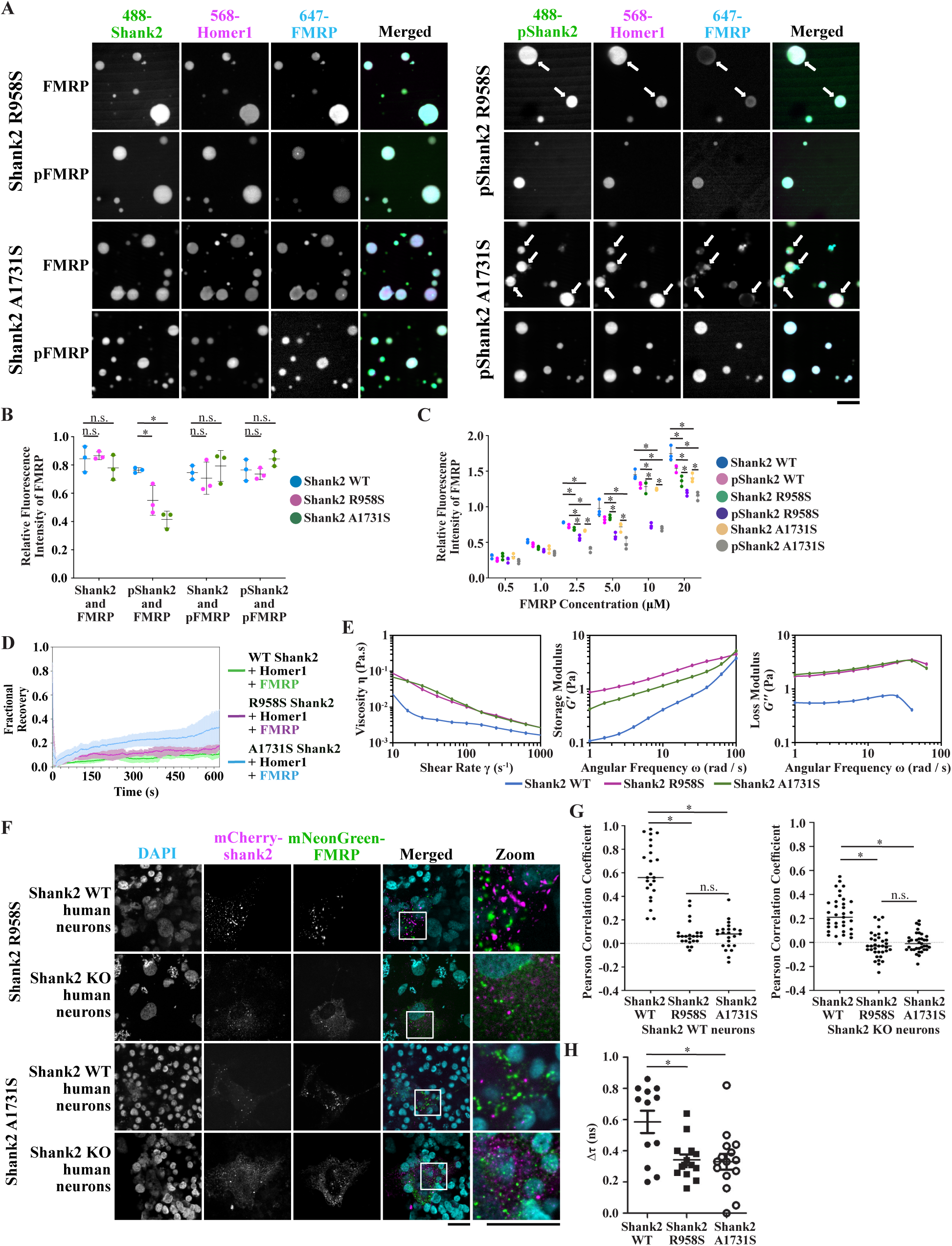
Disease-linked shank2 mutations promote compositional changes in condensates. **(A)** Confocal fluorescence microscopy images of AlexaFluor (AF) 488-R958S or A1731S (p)shank2 (green), AF568-homer1 (magenta), and AF647-(p)FMRP. White arrows highlight lack of FMRP-enrichment in R958S and A1731S pshank2 condensates compared with other conditions. FMRP enrichment occurs mostly on the edges of the shank2 / homer1 condensate. Scale bar = 10 μm. N = 3 experiments with 5 fields of view each analyzed. **(B)** Quantification of images shown in **(A)**. Intensity of AF647-(p)FMRP was divided by intensity of AF488-(p)shank2 to normalize data across conditions. Blue = WT shank2, Magenta = R958S shank2, Green = A1731S shank2. n.s. = not significant; * = p < 0.05, t-test. **(C)** Quantification of relative FMRP enrichment in shank2 (0.5 μM) and homer1 (0.5 μM) condensates when FMRP is increased from 0.5 μM to 20 μM. Blue = WT shank2, Pink = WT pshank2 WT, Green = R958S shank2, Violet = R958S pshank2, Gold = A1731S shank2, Silver = A1731S pshank2. * = p < 0.05, t-test. N = 3 experiments with 5 fields of view each analyzed. **(D)** Fluorescence recovery after photobeaching curves of FMRP shows the time course of recovery for FMRP in condensates formed with WT, R958S, or A1731S pshank2, homer1, and FMRP. Shown are the mean (green, magenta, cyan lines) ± s.d. (light green, light magenta, light cyan region above and below line). N = 3. **(E)** (left) WT (blue), R958 (magenta), and A1731S (green) pshank2, homer1, and FMRP condensates undergo shear thinning when increasing shear rates are applied. (center) Storage Modulus (G’) for WT (blue), R958S (magenta), and A1731S (green) pshank2, homer, and FMRP as angular frequency is increased. (right) Loss Modulus (G’’) for WT (blue), R958S (magenta), and A1731S (green) pshank2, homer, and FMRP as angular frequency is increased. Tables S1, S2, and S3 contain average and standard deviation for each condition. N = 3. **(F)** Confocal fluorescence microscopy images of 4-week-old SMAD+ NPC-differentiated inactive shank2 wild-type (WT) or shank2 knock out (KO) human neurons transfected with mCherry-R958S or A1731S shank2 (magenta) and mNeonGreen-FMRP (green). Zoomed images are of white box regions of merged images. Scale Bar under merged images = 20 μm. Scale bar under zoomed images = 5 μm. N = 6 experiments totaling 25-30 for each condition. **(G)** Pearson correlation coefficients measuring the colocalization of mNeonGreen-FMRP with mCherry-WT, R958S, or A1731S shank2 in NPC-differentiated human neurons shown in **(D)**. n.s. = not significant; * = p < 0.05. N = 6 experiments totaling 25-30 for each condition. **(H)** Fluorescence lifetime measurements from CHO cells expressing mNeonGreen FMRP and mCherry-WT, R958S, or A1731S shank2. Δτ is lifetime decay of mNeonGreen-FMRP from areas of mCherry-shank2 condensates and indicates strength of interaction between bulk mNeonGreen-FMRP and mCherry-WT, R958S, or A1731S shank2. * = p < 0.05, 1-way ANOVA.

One potential explanation for this observation is that mutant condensates have increased elasticity compared to WT condensates, as previously observed for WT and mutant TAF15 hydrogels (*21*). In presynaptic neuron terminals, the physical properties of condensates directly regulate the function of the active zone (*22*). Together, our results and previous observations from analogous systems led us to hypothesize that phosphorylation of R958S and A1731S shank2 mutants is physically more solid-like than those composed of WT shank2. We used oscillation rheology to probe the physical properties of WT and R958S or A1731S mutant pshank2 condensates. Probing of the viscosity of WT, R958S, or A1731S pshank2, homer1, and FMRP condensates under increasing shear rate resulted in decreasing viscosity, also called shear thinning, that is expected of non-Newtonian materials, like condensates (Fig. 3E). We observed that the storage and loss moduli of R958S and A1731S pshank2-containing condensates were increased compared to WT pshank2-containing condensates (Fig. 3E; tables S1, S2, S3), indicating that mutant pshank2, homer1, and FMRP condensates were indeed more elastic than their WT counterparts.

Expression of mCherry-R958S or A1731S shank2 with mNeonGreen-FMRP in WT or *SHANK2* KO human neurons at 4 weeks of differentiation resulted in the formation of distinct condensates of mutant shank2 or FMRP which were not colocalized with one another (Figs. 3F, G). This stands in contrast with significant colocalization between WT mCherry-shank2 and mNeonGreen-FMRP (Figs. 2E, 3G). These results provide evidence that heterozygous mutations, like those found in patients with one WT and one mutant copy of *SHANK2*, are dominant in a WT neuron. We then asked whether the strength of interaction between shank2 and FMRP was altered by R958S and A1731S mutations. Technical limitations for biochemical methods such as surface plasmon resonance prevented us from directly evaluating the strength of shank2 and FMRP interactions. Therefore, we used Fluorescence Lifetime Imaging Microscopy (FLIM) to probe shank2 and FMRP interactions in CHO cells. We found that these mutations result in significantly decreased interactions between shank2 and FMRP (Fig. 3H).

### Disease-linked mutation in shank2 result in altered mRNA translation

Control of condensate composition is essential for function (*23*, *24*). We hypothesized that lack of FMRP enrichment in shank2 condensates may result in an increase in RNA translation. We performed *in vitro* translation assays using rabbit reticulocyte lysates and luciferase mRNA that have previously been used to evaluate FMRP-dependent translation (*17*, *25*). We first asked whether lysates would alter 1) the formation of shank2 / homer1 condensates or 2) FMRP enrichment in shank2 / homer1 condensates. We observed no significant changes in condensate formation or FMRP enrichment in lysates compared with our previous solution-based assays (fig. S12 vs. Fig. 3A). We then performed *in vitro* translation assays using previously tested experimental conditions for FMRP enrichment in shank2 / homer1 condensates. Based on our model, we predicted that samples containing R958S or A1731S pshank2 would contain increased levels of luciferase due to higher levels of FMRP available for promoting RNA translation. As predicted, luciferase activity was significantly increased ∼15-fold and ∼30-fold in solutions containing R958S or A1731S pshank2, respectively (Fig. 4A). To this point, we have investigated only protein localization to shank2 condensates. Because condensates can enrich both protein and nucleic acids (*23*, *26*), we asked where luciferase mRNA localized in our assays. We observed that Cy5-labeled luciferase mRNA localized to WT shank2 condensates and was excluded from R958S or A1731S pshank2 condensates, similar to the observed pattern of FMRP enrichment (Fig. 4B, C).

**Fig. 4.**
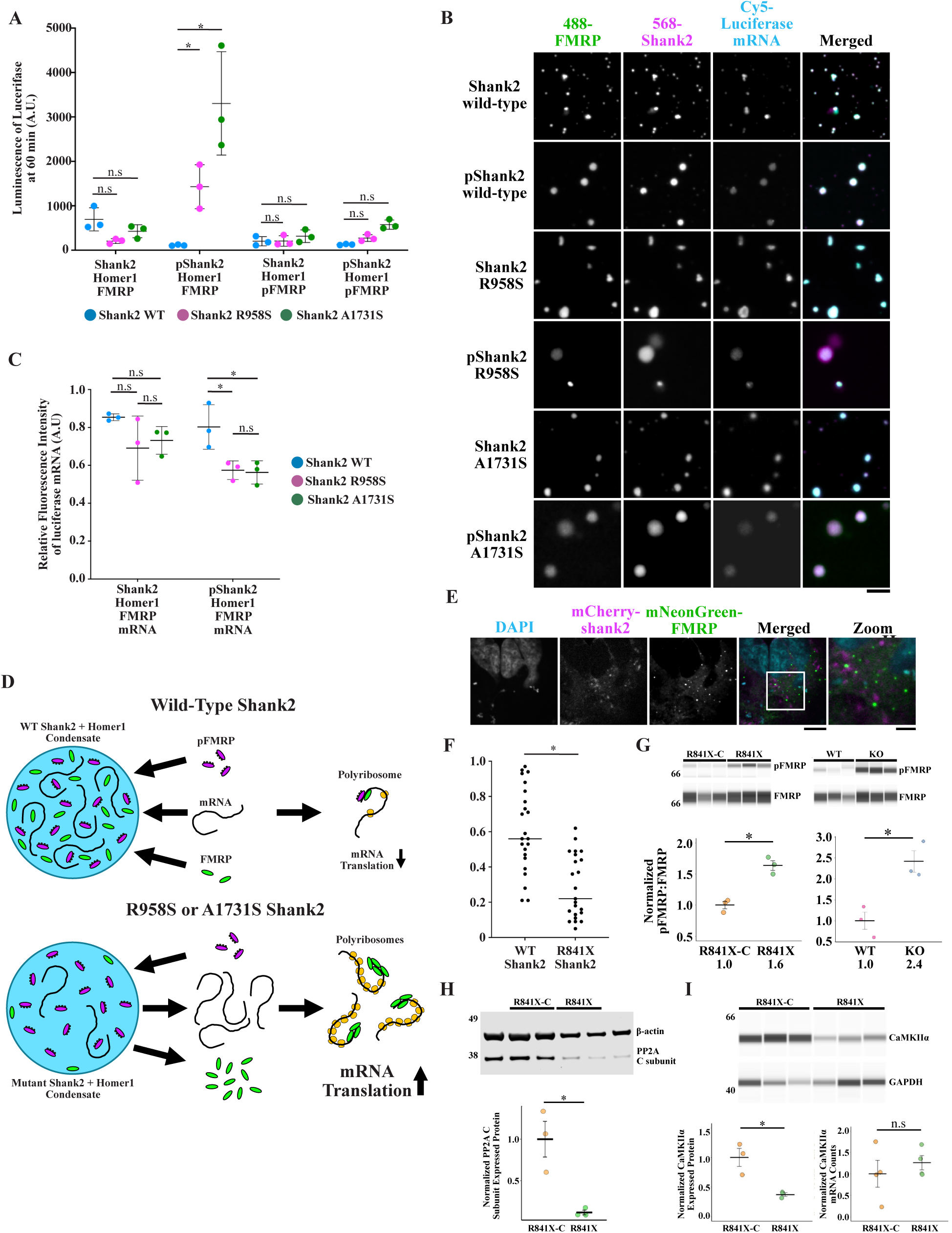
Mutations in shank2 result in aberrant RNA translation. **(A)** *In vitro* translation of luciferase mRNA in rabbit reticulocyte lysate after 60 minutes. All mixtures contain luciferase mRNA. Blue = WT shank2, Magenta = R958S shank2, Green = A1731S shank2. Shown are mean ± s.d. N = 3. * is p < 0.05, t-test. **(B)** Confocal fluorescence microscopy images of AlexaFluor (AF) 488-WT, R958S, or A1731S (p)shank2 (green), AF568-FMRP (magenta), and Cy5-luciferase mRNA. Scale bar = 5 μm. N = 3 experiments with 5 fields of view each analyzed. **(C)** Quantification of relative mRNA enrichment in shank2 (0.5 μM) and homer1 (0.5 μM) condensates with FMRP. Blue = WT shank2, Pink = WT pshank2 WT, Green = R958S shank2, Violet = R958S pshank2, Gold = A1731S shank2, Silver = A1731S pshank2. * = p < 0.05, t-test. N = 3 experiments with 5 fields of view each analyzed. **(D)** (top) Model of shank2-controlled mRNA translation. WT shank2 and homer1 condensates are enriched by FMRP, pFMRP, and mRNA resulting in dampened translation. (bottom) R958S and A1731S pshank2 and homer1 condensates are enriched by pFMRP, but FMRP and mRNA are not well-enriched resulting in increased translation compared to WT. **(E)** Confocal fluorescence microscopy images of 4-week-old *SHANK2* R841X shank2 heterozygous human neurons transfected with mCherry-WT shank2 (magenta) and mNeonGreen-FMRP (green). Zoomed images are of white box regions of merged images. Scale Bar under merged images = 20 μm. Scale bar under zoomed images = 5 μm. **(F)** Pearson correlation coefficients measuring the colocalization of mNeonGreen-FMRP with mCherry-WT shank2 expressed in WT or R841X shank2 NPC-differentiated human neurons shown in **(E)**. WT data is same as those shown in Fig. 3G. * = p < 0.05. N = 6 experiments totaling 25-30 for each condition. **(G)** Western blots of FMRP and pFMRP from isogenic pairs of *SHANK2* human neurons at 9 weeks of differentiation (R841X-C and heterozygous R841X; WT and homozygous KO). Intensity of FMRP and pFMRP bands was measured and normalized to FMRP. Ratio of intensities indicated under plots. * = p < 0.05, t-test. N = 3. **(H)** Western blot of PP2A subunit C protein expression in heterozygous R841X and corrected R841X (R841X-C) shank2 human neurons at 9 weeks of differentiation. Intensity of PP2A subunit C and β-actin bands was measured and PP2A subunit C intensity was normalized to β-actin. Ratio of intensities is indicated under plot. * = p < 0.05, t-test. N = 3 **(I)** Western blot of CaMKIIα protein expression in heterozygous R841X and corrected R841X (R841X-C) shank2 human neurons at 9 weeks of differentiation. Intensity of CaMKIIα and GAPDH bands was measured and CaMKIIα intensity was normalized to GAPDH. Ratio of intensities is indicated under plots. CaMKIIα mRNA counts in R841X and R841X-C *SHANK2* human neurons at 9 weeks of differentiation detected by RNA-Seq (*8*). n.s. = not significant, * = p < 0.05, t-test. N = 3

Collectively, these results enabled us to develop a novel mechanistic model in which shank2 indirectly modulates mRNA translation by regulating the enrichment and sequestration of FMRP, pFMRP, and mRNA (Fig. 4D). Any RNA outside of shank2 condensates is free to be translated when bound by FMRP but repressed when bound to pFMRP (*17*, *25*). When R958S or A1731S shank2 is phosphorylated, FMRP and RNA are not enriched like in WT condensates (Fig. 2B vs. 3A, 4C), while pFMRP is highly enriched (Fig. 3A), thereby shifting the ratio of FMRP to pFMRP and levels of mRNA outside of shank2 condensates, underlying the observed increase in translation.

To this point, we have discovered that shank2 mutations can directly increase the elasticity and alter the composition of condensates, resulting in increased RNA translation (Fig. 4D). To examine shank2-FMRP pathway alterations, we used iPSC neuron lines WT, *SHANK2* homozygous KO, heterozygous C-terminal IDR deletion R841X *SHANK2* that expresses half the WT shank2 levels, and rescued R841X-C *SHANK2* in which CRISPR was used to fuse the C- terminus of *SHANK2* with the first 841 residues of the truncated mutant. We first assessed the co-localization of mNeonGreen-FMRP and mCherry-shank2 in condensates in R841X *SHANK2* neurons and found that, like R958S and A1731S shank2 mutants, FMRP co-localizes poorly with shank2 (Fig. 4E, F). We probed pFMRP and FMRP levels in WT and KO *SHANK2* neurons and found that the ratio of pFMRP:FMRP is significantly increased in KO vs. WT (Fig. 4G).

Likewise, pFMRP:FMRP was significantly higher in R841X *SHANK2* vs. R841X-C neurons (Fig. 4G). PP2A is a known phosphatase of FMRP, so we asked whether PP2A levels are altered in R841X neurons compared with rescued R841X-C neurons. Consistent with increased pFMRP:FMRP ratios, we found that the expression of the C subunit of PP2A is significantly decreased in the R841X neurons vs. rescued neurons (Fig. 4H). Together, these data indicate that in R841X mutants, mislocalization of FMRP away from shank2 condensates results in genetic perturbations that lead to decreased phosphatase expression and an increased pFMRP:FMRP ratio. Because pFMRP can repress RNA translation (*25*), we predicted that an increased pFMRP:FMRP ratio will result in decreased expression levels of FMRP targeted RNA. We therefore probed the expression of CaMKIIα, an essential neuronal protein that is under the translational control of FMRP (*27*), in active neurons at 9-weeks post differentiation. Indeed, CaMKIIα expression in R841X neurons is decreased compared to rescued R841X-C shank2 neurons even though RNA counts in cells are similar (Fig. 4I). Overall, our findings support a clear connection between shank2 and FMRP-modulated translation of neuronal proteins

## Summary

In summary, our results provide the first evidence of a direct link between biomolecular condensates containing shank2 and FMRP modulated mRNA translation. We identified two ‘classes’ of shank2 mutations that result in dysregulated FMRP-modulated translation: 1) serine missense mutations that lead to reduced enrichment of FMRP in shank2 condensates resulting in increased mRNA translation, and 2) null mutations that result in reduced FMRP enrichment in shank2 condensates coupled with an increased pFMRP:FMRP ratio that led to decreased mRNA translation. Condensates that are composed of mutant shank2 and homer1 effectively act as a selective filter with respect to FMRP: FMRP is selected against and not enriched in the condensates while pFMRP is selected for and is well enriched. These observations have major implications for understanding vast numbers of disease-linked missense mutations found within condensate-associated proteins and suggest that, like ALS-associated protein mutations (*28*, *29*), unique mutations can share molecular phenotypes. Our cellular results show that these missense mutations are dominant in WT background human neurons and form condensates in the absence of endogenous shank2 in KO neurons. Perhaps most strikingly, our data exemplify a link between single missense mutations, increased physical elasticity of mutant condensates and significantly altered composition, and dysregulated functional outputs. Collectively, these results establish a new framework through which to investigate condensate-related diseases.

## Acknowledgements

We thank Dr. Greg Wasney and the Structural & Biophysical Core at SickKids Research Institute for help with instruments used in RNA translation experiments and rheometry, the SickKids Imaging Facility for help with microscopy, Dr. Hyun O Lee for help with FRAP, Drs. Julie Forman-Kay and Rhea Hudson for MAPK protein and FMRP plasmid, and Dr. Roman Melnyk and members of the Ditlev, Lee, and Ellis Labs for helpful discussions. This work was supported by funds from the SickKids Research Institute (to J.A.D.); Simons Foundation Autism Research Initiative (Grant No. 514918 to J.E.), the Ontario Brain Institute (POND Network to JE); John Evans Leaders Fund/Canada Foundation for Innovation (JELF/CFI) and Ontario Research Fund (to J.E.), Canada Research Chair (Tier 1) in Stem Cell Models of Childhood Disease (to J.E.), the Beta Sigma Phi International Endowment Fund (to J.E.); Ontario Graduate Student Award (to F.M.P.), David Stephen Cant Scholarship in Stem Cell Research (to F.M.P.); Canadian Institute of Health Research project grants (PJT-156034 and PJT-156439 to L.-Y.W.), and Canada Research Chair (Tier 1) in Brain Development and Disorders (to L.-Y.W.).

## Data Availability

All original data will be uploaded to an online data repository upon acceptance of the manuscript for publication.

## Supplemental Materials

Methods and Materials Figures S1–S12

Table S1-S4

Author Contributions

Supplemental References (*30–37*)

