## Supplemental Material for "Disease-linked mutations dysregulate neuronal condensate physical properties, composition, and RNA translation"

### Supplementary Materials for

**Title: Disease-linked mutations dysregulate neuronal condensate physical properties, compositional control, and RNA translation**

**Authors:** Leshani Ahangama Liyanage, Fraser McCready, Steve Chung, Jason Arsenault, Wei Wei, Xusheng Lin, Lu-Yang Wang, James Ellis, Jonathon A. Ditlev

**The PDF file includes:**

Materials and Methods
Figs. S1 to S12
Tables S1-S4
Author Contributions
Supplemental References (*30-37*)

#### Materials and Methods

Plasmid generation

The gene encoding full-length human SHANK2a was codon optimized and synthesized by GenScript bound by EcoRI and NotI restriction enzyme cloning sites. *SHANK2a* was transferred to a pSUMO vector to generate an N-terminal His-tagged SUMO fusion protein for expression in *Escherichia coli*. All *SHANK2a* mutants (R958S, G1172R, R1429W, and A1731S) were generated by the standard PCR-based method using Phusion Site-directed Mutagenesis kit (Thermo Fisher Scientific). The gene encoding *HOMER1* was purchased from the UT Southwestern Sanger Sequencing Core Ultimate ORF Lite human cDNA collection (Thermo Fisher Scientific). *HOMER1* was amplified by PCR using forward primer 5'-CCCCCGGATCCGGGAACAACCTATCTTCAGC and reverse primer 5'-CCCCCGCGGCCGCTCATTACTAGCTGCATTCTAGTAGCTTGGCCAA. Following BamHI and NotI digestion, HOMER1 was ligated into a BamHI / NotI digested pSUMO vector. The pET SUMO plasmid encoding *FMRP* residues 445 - 632 was generously provided by the Forman-Kay Lab (*30-31*). Full-length *SHANK2*, *SHANK2* mutants, and *FMRP* were cloned into pmCherry-N1 and pmNeonGreen-N1 vectors for mammalian cell expression under a CMV promoter. *SHANK2* and *SHANK2* mutants were amplified by PCR using forward primer 5'-CCAGTGAATTCATGAAGAGCC-3' and reverse primer 5'-GCGGCCGCGGATCCGTACGATCCAGAAGCTGC-3'. Following digestion with EcoRI and BamHI, *SHANK2* or *SHANK2* mutants were ligated into a EcoRI / BamHI digested pmCherry-N1 vector. *FMRP* was digested from a parent *FMRP*-CFP plasmid (*31*) using XhoI and XmaI and ligated with a XhoI / XmaI digested pmNeonGreen-N1 vector. All constructs were confirmed by Sanger sequencing. mCherry-shank2 and mCherry-shank2 mutant or FMRP-mNeonGreen expression in CHO cells was assessed by western blot using anti-shank2 antibody (Cell Signaling, 12218) or anti-FMRP antibody (Biolegend, 834701), respectively.

Protein Expression and Purification

*Shank2*

His-SUMO-shank2 wild type and / or mutants were transformed into Rosetta-2 (DE3) cells in LB and grown at 37 °C. Protein expression was induced with 0.5 mM IPTG when the culture grew to $O.D._{600nm}$ ~0.6 and incubated at 16 °C for 18 hours followed by 4 °C. Cells were harvested and pelleted via centrifugation at 4500 rpm for 30 minutes at 4 °C (Thermo Fisher Scientific, Sorvall Lynx 6000). Cell pellets were resuspended in lysis buffer containing 6M guanidine hydrochloride (GdnHCl), 20 mM imidazole (pH 8), 150 mM NaCl, 10% glycerol, 0.01% NP-40, 5 mM β-mercaptoethanol (βME), 1 µg / ml antipain, 1 µg / ml pepstatin, 1 µg / ml leupeptin and 0.1 mM phenylmethane sulphonyl fluoride (PMSF). Cells were lysed and homogenized using cell homogenizer (Emulsiflex-C3, Avestin) at 10,000 psi, 3X at 4 °C and lysates were collected ice. Lysates were cleared by centrifugation at 14,500 rpm for 1 hour at 4 °C and loaded onto Ni-NTA agarose resin (Invitrogen) equilibrated with lysis buffer. The Ni-NTA agarose resin was washed with 5 column volumes of wash buffer 1 containing 50 mM Tris (pH 8), 150 mM NaCl, 5 mM βME, 0.01 % NP-40, 10 % (Volume:Volume) glycerol ("Buffer A") with 3 M GdnHCl. Then resin was washed 5 column volumes of wash buffer 2 containing Buffer A and 1.5 M GdnHCl, followed by 5 column volumes of wash buffer 3 containing Buffer A and 0.75 M GdnHCl. Elution buffer containing 50 mM Tris (pH 8), 300 mM imidazole (pH 8), 150 mM NaCl, 5 mM βME, 0.01 % NP-40, 10 % glycerol and 0.75 M GdnHCl was used to elute His-SUMO-shank2. SDS-PAGE gel was used to confirm the elution fractions containing shank2.

Elution fractions were concentrated by Amicon Ultra Centrifugal Filter 50 k units (EMD Millipore) and further purified by size exclusion chromatography using HiLoad 16 / 600 Superose 6 column (Cytiva) equilibrated with buffer containing 25 mM HEPES (pH 7), 300 mM NaCl, 0.75 M GdnHCl, 150 mM Arginine, 10 % glycerol and 1 mM (Tris(2-carboxyethyl)phosphine hydrochloride (TCEP). SDS-PAGE was used to identify shank2 containing fractions and they were pooled together and concentrated. Single-use aliquots were flash frozen in liquid nitrogen and stored at -80 °C for future use. Cleaving the SUMO-fusion tag before size-exclusion column resulted in partial loss of shank2. Thus, to simplify the procedure and to obtain a good yield, SUMO tag was retained during purification and condensate formation was initiated by ULP-1 cleavage. Residue sequence can be found in table S4.

*Homer1*

The gene encoding human full-length Homer1 protein was obtained from the Ultimate ORF clone library (Invitrogen). The fusion protein was designed as pSUMO vector with a N-terminal $His_6$-SUMO tag for expression in *Escherichia coli*. BL21(DE3) cells containing $His_6$-SUMO-homer1 were grown in LB at 37 °C. Protein expression was induced with 0.5 mM IPTG when the culture grew to $O.D._{600nm}$ ~0.6 and incubated at 16 °C for 18 hours followed by 4 °C. Cells were collected by centrifugation at 4500 rpm for 30 minutes at 4 °C (Thermo Fisher Scientific, Sorvall Lynx 6000) and lysed by cell disruption (Emulsiflex-C3, Avestin) at 10,000 psi, 3X at 4 °C in 10 mM imidazole (pH 8), 150 mM NaCl, 5 mM β-ME, 0.01% NP-40, 10% glycerol, 0.1 mM PMSF, 1 μg / ml antipain, 1 μg / ml pepstatin, and 1 μg / ml leupeptin. Lysates were collected on ice. Lysates were cleared by centrifugation at 14,500 rpm for 1 hour at 4 °C. Centrifuge-cleared lysate was applied to Ni-NTA agarose resin (Invitrogen), washed first with 20 mM imidazole (pH 8), 500 mM NaCl, 5 mM βME, and 10% glycerol, then washed with 50 mM imidazole (pH 8), 250 mM NaCl, 5 mM βME, and 10% glycerol, and eluted with 300 mM imidazole (pH 8), 250 mM NaCl, 5 mM βME, and 10% glycerol. His-SUMO tag was cleaved by ULP-1 protease treatment for 16 hours at 4 °C or for 2 hours at room temperature. SDS-PAGE gel was used to confirm successful cleavage of the His-SUMO tag. Cleaved homer1 was applied to a Source 15Q anion exchange column (Cytiva) and eluted with a gradient of 0 to 1M NaCl in 20 mM HEPES (pH 7), 1 mM DTT, 10% Glycerol. Homer1 was further purified by size exclusion chromatography using a HiLoad Superdex 200 prepgrade column (Cytiva) in 20 mM Tris-HCl (pH 8), 500 mM NaCl, 5 mM βME and 10% glycerol. Elution fractions were concentrated by Amicon Ultra Centrifugal 10k filter units (EMD Millipore). Residue sequence can be found in table S4.

*FMRP (445-632)*

Expression plasmid for the low-complexity disordered c-terminal of human FMRP (445-632) was a generous gift from the Forman-Kay Lab. The methodology for expressing and purifying FMRP (445-632) was adapted from a previously described approach (*30,31*). Briefly, His-SUMO-FMRP was transformed into *Escherichia coli* BL21(DE3) cells and grown at 37 °C in LB. Protein expression was induced with 0.5 mM IPTG when the culture grew to $O.D._{600nm}$ ~0.6 and incubated at 18 hours at 16 °C followed by 4 °C. Cells were harvested and pelleted via centrifugation at 4500 rpm for 30 minutes at 4 °C (Thermo Fisher Scientific, Sorvall Lynx 6000). Cells were lysed by cell disruption (Emulsiflex-C3, Avestin) at 10,000 psi, 3X at 4 °C in buffer containing 6 M GdnHCl, 50 mM Tris (pH 8), 500 mM NaCl, 20 mM imidazole, and 2 mM βME, and 0.1 mM PMSF, 1 μg / ml antipain, 1 μg / ml pepstatin, and 1 μg / ml leupeptin. Lysates were collected on ice. Lysates were cleared by centrifugation at 14,500 rpm for 1 hour at 4 °C and the lysate was loaded onto Ni-NTA agarose resin (Invitrogen) equilibrated with lysis buffer. The Ni-NTA resin was washed with 10 column volumes of lysis buffer followed by 5 column volumes of lysis buffer without 6 M GdnHCl. Elution buffer containing 50 mM Tris pH 8, 500 mM NaCl, 300 mM imidazole, and 2 mM βME was used to elute His-SUMO-FMRP. The His-SUMO tag was cleaved with ULP-1 protease while dialyzed against cleavage buffer containing 50 mM Tris pH 8, 150 mM NaCl, 20 mM imidazole, 2mM βME overnight. FMRP was separated from the His-SUMO tag by loading the protein onto Ni-NTA agarose resin and collecting the flow-through. SDS-PAGE gel was used to confirm successful cleavage of the His-SUMO tag and separation of the tag from FMRP. The flow-through was concentrated using Amicon Ultra Centrifugal 3k filter units and further purified using a HiLoad Superdex 75 26 / 600 size-exclusion column (Cystiva) equilibrated with buffer containing 4 M GdnHCl, 50 mM Tris pH 8, 500 mM NaCl, and 2 mM βME. FMRP containing fractions were pooled together. The purity of FMRP was confirmed with SDS-PAGE. Residue sequence can be found in table S4.

<u>Fluorophore Conjugation</u>

*Shank2*

A fluorescence dye was conjugated via maleimide linkage to any of nine cysteine residues present in wild-type (WT) and mutant shank2. Purified WT or mutant shank2 were concentrated using Amicon Ultra Centrifugal 50k filter units (EMD Millipore). For the labeling reaction, either Alexa Fluor (AF) 488 C5 Maleimide (Invitrogen) or AF568 C5 Maleimide (Invitrogen) was added in protein: dye ratio of 1: 2 to WT or mutant shank2 in 25 mM HEPES (pH 7.0), 300 mM NaCl, 0.75 M GdnHCl, 150 mM Arginine, 10% glycerol, and 1 mM TCEP. The reaction was incubated with gentle mixing at 4 °C for 16 hours. The reaction was quenched with 1 µl 14.3 M βME the following day. The fluorophores and other small molecules were removed from the protein by size exclusion chromatography using a Superose 6 Increase 10 / 300 GL column (Cytiva) with buffer containing 25 mM HEPES (pH 7), 300 mM NaCl, 0.75 M GdnHCl, 150 mM Arginine, 10% glycerol and 1 mM TCEP. Fractions containing fluorescent-labeled WT or mutant shank2 were pooled and concentrated using Amicon Ultra Centrifugal Fiter units (Millipore). Successful dye separation was confirmed by SDS-PAGE gel and labeled protein concentration was determined by $A_{280}$ and $A_{488}$ / $A_{568}$ absorption, depending on the AF maleimide dye, using a Nanodrop One-C (Thermo Fisher Scientific).

*Homer1*

A fluorescence dye was conjugated using a maleimide linkage to two cysteine residues located at amino acid positions 234 and 353 within homer1. Purified homer1 was concentrated using Amicon Ultra Centrifugal 10k filter units (EMD Millipore) to ~ 100 µM. βME was added to 5 mM to reduce cysteine residues followed by buffer exchange using a HiTrap 26 / 10 Desalting column (Cytiva) in 25 mM HEPES (pH 7) and 150 mM NaCl to ensure that any residual reducing agents were removed. Fractions containing protein were collected and concentrated to 100 µM. 200 µM AF568 C5 Maleimide (Invitrogen) was added, and the reaction was incubated with gentle mixing at 4 °C for 16 hr. The reaction was quenched with 1 µl 14.3 M βME the following day and unreacted dye was removed by buffer exchange using the desalting column previously described in 25 mM HEPES (pH 7), 150 mM NaCl, and 1 mM βME followed by size exclusion chromatography using a Superdex 200 Increase 10 / 300 GL column (Cytiva) in 25 mM HEPES (pH 7), 150 mM NaCl, and 1 mM βME. Fractions containing AF568-labeled homer1 were pooled and concentrated using Amicon Ultra Centrifugal 10k filter units (EMD Millipore), confirmed the presence of labeled protein by SDS-PAGE and protein concentration was determined by $A_{280}$ and $A_{568}$ absorption, using a Nanodrop One-C (Thermo Scientific).

*FMRP*
A fluorescence dye was conjugated to a single cysteine residue in FMRP via maleimide linkage. For labeling reaction purified FMRP was concentrated using Amicon Ultra Centrifugal 3k Filter units (Millipore) to ~ 100 µM. βME was added to 5 mM to reduce cysteine residues followed by buffer exchange using a HiTrap 26 / 10 desalting column (Cytiva) in 25 mM HEPES (pH 7.5) and 4M GdnHCl to ensure that any residual reducing agents were removed. The fractions containing protein were collected and concentrated to 100 µM. Then immediately reacted with either AF488 C5 Maleimide (Invitrogen) or AF647 C2 Maleimide (Invitrogen) at protein: dye ratio of 1: 0.1 or 1: 1, respectively. The reaction was incubated with gentle mixing at 4 °C for 16 hours. The reaction was quenched with 1 µl 14.3 M βME on the following day. Unreacted dye was removed by buffer exchange using the desalting column described above in 25 mM HEPES (pH 7.5), 4 M GdnHCl, and 1 mM βME followed by size-exclusion chromatography using a Superdex 75 column Increase 10 / 300 GL column (Cytiva) in 25 mM HEPES (pH 7.5), 4M GdnHCl, and 1 mM βME. Fractions containing labeled FMRP were pooled and concentrated using Amicon Ultra Centrifugal 50k filter units (EMD Millipore). Successful dye separation was confirmed by SDS-PAGE gel analysis and labeled protein concentration was determined by $A_{280}$ and $A_{488}$ / $A_{647}$ absorption, depending on the AF maleimide dye, using a Nanodrop One-C (Thermo Fisher Scientific). Final protein concentrations of all labeled proteins and degree of labeling were calculated from the protein absorbance using the following formulas:

AF488
Concentration (M) = A280 – (A495x0.11)/ ExtCoef
Degree of labeling = A495/(71,000x [protein conc.])

AF568
Concentration (M) = A280 – (A568 x 0.46)/ExtCoef
Degree of labeling = A568/(92,000x [protein conc.])

AF647
Concentration (M) = A280 – (A650 x 0.03)/ExtCoef
Degree of labeling = A650/(239,000x [protein conc.])

Protein Phosphorylation Reactions
WT or mutant shank2 were phosphorylated using Mitogen-activated protein (MAP) kinase, while FMRP was phosphorylated using Casein kinase II (New England Biolabs). MAP kinase was received as a generous gift from Rhea Hudson of the Forman-Kay Lab. Both purified proteins were added separately to phosphorylation reaction buffer containing 25 mM HEPES (pH 7.5), 2 mM DTT, 150 mM NaCl, 20 mM $MgCl_2$ and 15 mM ATP. 10 ng / µl of MAP kinase or CK2 was added to the corresponding protein and the reaction was incubated in 30 °C for 24 hours (*32-35*). Due to challenges in handling WT and mutant shank2, the phosphorylation reaction mixtures were kept without further purification. The phosphorylation of shank2 proteins

were confirmed via mass spectrometry. Phosphorylation of FMRP was confirmed using SDS-PAGE gel analysis.

Reconstitution Assays

Prior to use, 384-well glass-bottomed plates (Corning) were washed with Hellmanex III (Hëlma Analytics) for 3.5 hours at 50 °C and thoroughly rinsed with MilliQ $H_2O$. Individual wells were washed with 6 M NaOH for 30 minutes at 45 °C two times to strip the exposed layers of glass, thoroughly rinsed with MilliQ $H_2O$ three times, followed by equilibration with 50 mM HEPES (pH 7.3), 150 mM NaCl, and 1 mM TCEP ("Buffer B"). The wells were blocked with Buffer B containing 10 mg / ml BSA for 30 minutes at room temperature and Buffer B was replaced with protein solutions. Purified WT or mutant shank2, homer1, FMRP and phosphorylated proteins of varying concentrations were prepared in buffer containing 50 mM HEPES (pH 7.5), 1 mM TCEP and 1 mg / ml BSA, and immediately placed into wells in the prepared 384-well plate. Unless otherwise noted, when fluorescently labeled proteins were mixed at a 20 % labeled protein and 80 % unlabeled protein ratio to avoid photochemical effects (*36*). The plate was incubated for 30 minutes at room temperature prior to imaging.

FMRP Enrichment in Shank2 and Homer1 Condensate Across Concentrations

Overall, we sought to use physiologically relevant concentrations of shank2, homer1 and FMRP in the *in vitro* reconstitution reactions. To assess the enrichment of FMRP within shank2 and homer1 condensates over a concentration range, we maintained a constant shank2 and homer1 concentrations of 0.5 μM while varying the FMRP concentration from 0.5 μM to 20 μM. We employed confocal imaging of condensates, using *in vitro* reconstitution in solution for this analysis. Briefly, purified non-phosphorylated and phosphorylated WT or mutant shank2, homer1 and FMRP were prepared at the desired concentrations in buffer containing 50 mM HEPES (pH 7.5), 1 mM TCEP and 1 mg / ml BSA. First shank2 and homer1 were loaded into the wells of a prepared imaging plate, and after 30 minutes, FMRP was added. The plate was then incubated for 10-20 minutes and imaged for condensate formation. Confocal images were captured using Leica DMi8 spinning disk microscope equipped with a Hamamatsu C9100-13 EM-CCD camera with 63x / 1.4 oil objective. Images were analyzed using FIJI image analysis software.

Imaging of Biomolecular Condensates Using Spinning Disk Confocal Microscopy

Confocal images of condensates were captured using Leica DMi8 spinning disk microscope equipped with a Hamamatsu C9100-13 EM-CCD camera with 63x / 1.4 oil objective. Alexa fluor 488, 568 and 647 fluorescence were detected using a 488 nm (150 mW), 561 nm (150 mW) and 637 nm (140 mW) laser respectively. Images represent droplets settled to the bottom of the plate. Images were analyzed using Volocity (Perkin Elmer) and FIJI image analysis software.

Fluorescence Recovery After Photobleaching (FRAP)

FRAP experiments were performed on a Nikon Ti-2E confocal microscope equipped with an Andor Dragonfly 200 unit, 60x / 1.4 oil objective, and Andor Zyla sCMOS camera. Either Alexa Fluor (AF) 488 or AF647 labeled WT or mutants shank2, homer1 and either AF488 or AF647 labeled FMRP were mixed in buffer containing 50 mM HEPES (pH 7.5) and 1 mM TCEP to form droplets. Varying mixtures of phosphorylated and / or unphosphorylated WT or mutant shank2 and FMRP were used for FRAP experiments. Pre-mixed protein samples are placed on cleaned 384-well glass-bottomed plates (Corning) and incubated at room temperature (~5 minutes). A diameter of ~1 - 2 µm was bleached for 8 seconds with 100% laser power at 488 nm to photobleach AF488-labeled shank2 WT/mutants or FMRP. A single confocal plane was maintained, and images were collected for 10 minutes. The fluorescence intensity difference between pre-bleaching and at time 0 was normalized to 100%. Images were processed using Imaris Viewer (Oxford Instruments) and analyzed using FIJI image analysis software.

*In Vitro* Translation Assay

*In vitro* translation was performed using Luciferase mRNA (Promega) and the nuclease treated Rabbit Reticulocyte Lysate System (Promega). Each reaction (30 µL) contained 12.6 µL of rabbit reticulocyte lysate, 0.5 µL of Luciferase mRNA (1 mg/mL), 0.3 µL of amino acid mixture minus leucine (1 mM), 0.3 µL of amino acid mixture minus methionine (1 mM), and 16.3 µL of protein mixture or buffer (50 mM HEPES pH 7.5, 1 mM DTT, and nuclease-free $H_2O$). Protein components used in the assay include WT or mutant Shank2 or phosphorylated WT or mutant Shank2, Homer1, and FMRP or phosphorylated FMRP. Translation was reported as the increase in luminescence at the endpoint of 1 hour using a standard luciferase assay system (Promega). A standard reaction contained 75 µL of luciferase substrate mixed with 3 µL of unpurified translation mixture in a white opaque 96 well plate (Costar). Luminescence was measured using a Synergy Neo2 Multi-Mode Assay Microplate Reader (BioTek) at 25 °C. Relative luminescence measurements represent mean values obtained from three independent *in vitro* translation reactions, each of them measured two times.

Confirmation of Condensates in *In Vitro* Translation Assays

The presence of droplets in translation assay samples were observed using Leica DMi8 spinning disk microscope equipped with a Hamamatsu C9100-13 EM-CCD camera with 63x / 1.4 oil objective. Initially droplet formation was observed as Differential Interference Contrast (DIC) images, however, the background effect of Rabbit Reticulocyte Lysate resulted in difficulty of seeing droplets clearly. Thus, to observe the droplets clearly in translation assay samples, we repeated the microscopy assays described above with 20 % of fluorescent-labeled protein mixed with 80% unlabeled protein in translation assay Rabbit Reticulocyte Lysate (Promega) mixtures.

Localization of Cy5-labeled Luciferase mRNA in Condensates

To investigate the localization of mRNA within shank2, homer1, and FMRP condensates, we conducted an *in vitro* experiment using Cy5-labeled luciferase mRNA (APExBIO) along with fluorophore-labeled shank2 and FMRP. The concentrations for phosphorylated and non-phosphorylated shank2 WT or mutants and homer1 were set at 1 µM, while FMRP was at 5 µM. The concentration of luciferase mRNA was 0.0015 µM. *In vitro* reconstitution was performed as previously described, starting with the addition of shank2 and homer1, followed by the addition of FMRP and mRNA 30 minutes later in a buffer containing 50 mM HEPES (pH 7.5), 50 mM NaCl, 1 mM TCEP, and 1 mg/ml BSA. Images of shank2, homer1, FMRP and luciferase mRNA condensates were captured on Leica DMi8 spinning disk microscope equipped with a Hamamatsu C9100-13 EM-CCD camera with 63x / 1.4 oil objective. Images were analyzed using FIJI image analysis software.

Oscillation Rheology

pShank2, homer1, and FMRP were mixed to a concentration of 5 μM each in 50 mM HEPES (pH 7.5), 50 mM NaCl, and 1 mM TCEP and centrifuged at 500 g for 30 minutes at room temperature to pellet condensates. The Discover HR-3 Rheometer (Waters TA Instruments) was zeroed. Geometry, including friction, inertia, and rotational mapping, was calibrated and environmental temperature equilibrated at 25 °C. Condensate containing sample was loaded onto Peltier-controlled sample plate and the gap was trimmed to 200 μm and then 100 μm to prevent trapping air between the plate and sample. Sample was equilibrated for 1 minute to allow material to reach 25 °C. Shear rate experiments were performed from 10 $s^{-1}$ to 1000 $s^{-1}$ with an equilibrating time of 5 s and averaging time of 15 s. Oscillation experiments to measure Storage and Loss Moduli were performed with 50 % strain and 1 rad / s to 100 rads / s with an equilibrating time of 5 s and averaging time of 15 s.

<u>iPSC-derived neuron differentiation and culturing</u>

iPSCs were reprogrammed under approval of the Canadian Institutes of Health Research Stem Cell Oversight Committee and the Research Ethics Board of The Hospital for Sick Children (REB #1000050639). iPSC-derived Neural Progenitor Cells (NPC) were isolated and differentiated as described (*37*). The isogenic NPC cultures (wild-type and homozygous *SHANK2* knockout, or heterozygous *SHANK2* R841X and its corrected control R841X-C) were thawed at 37 °C and transferred to 10 ml of NPC media containing 1x DMEM F12, 100x N2 supplement, 100x NEAA, 100x penicillin/streptomycin, 50x B27, 1μg/ml laminin and 2μg/ml heparin. NPCs were spun at 1100 rpm for 5 minutes, supernatant was removed, and cells were resuspended in NPC media containing fresh 20ng/m FGF2. NPCs were then placed in matrigel coated 6-well plate and incubated at 37 °C for one week, changing NPC media with fresh FGF2 every other day.

For NPC seeding PORN / Laminin coated surfaces were prepared. Briefly, coverslips were placed in the middle of a 24-well plate, coated them with 0.1 mg/ml PORN diluted in RNAase/DNAase free $H_2O$ and placed overnight at 37 °C. Following day, the PORN was aspirated, and the plate was washed one time with RNAase / DNAase free $H_2O$. The plate was air dried with lid open for as long as needed (1~2 hours). To coat with laminin, 20μg /ml laminin was diluted in (1:25) DMEM F12 and 80μl of laminin drop was placed to middle of each coverslip. The plate was placed at 37 °C for 2 hours or overnight.

The NPC-containing 6-well plate was observed under the microscope and the old media was aspirated. The plate was washed with 1x PBS buffer and aspirated. NPCs were digested with Accutase for 5 minutes at 37 °C to lift them off the plate and carefully triturated using a P1000 pipette a maximum of 15 times, washed with NPC media and counted. Concomitantly, laminin was aspirated from PORN / laminin coated 24-well plate. The neuronal differentiation (CND) medium containing Neurobasal, 100x N2, 50x B27, 100x GlutaMAX-1, 100x penicillin/streptomycin supplemented with cAMP (1μM), BDNF (10ng /ml), GDNF (10ng /ml), ascorbic acid (200ng /ml), 1μg/ml laminin and 10μM rock inhibitor was added to each well. NPCs were seeded at a density of 30,000 per well and incubated at 37 °C. The cells were fed with CND medium twice a week by changing 75% of the medium in each well. For days 10–14, the cultures were treated with 5μM DAPT to attempt to curb excess NPC proliferation and promote differentiation.

<u>Imaging and Analysis of Neurons</u>

Fixed neurons were imaged with Leica DMi8 spinning disk microscope equipped with a Hamamatsu C9100-13 EM-CCD camera with 63x / 1.4 oil objective. DAPI, mCherry-shank2 wt or mutants and full length mNeonGreen-FMRP were visualized using 405 nm, 561 nm, and 488 nm lasers, respectively. Individual channels were acquired in sequential manner to prevent bleed-through of fluorophores. We aimed to acquire at least ten neurons per coverslip, acquired in separate batches. Images were analyzed using Volocity (Perkin Elmer) and FIJI image analysis software. The same brightness and contrast were applied to images within the same channels. Camera background was subtracted before calculating mean fluorescence intensities. After thresholding of signal-containing areas with the 'triangle' algorithm, a binary mask was created. mCherry-shank2 wt or mutants and mNeonGreen-FMRP puncta within neurons were analyzed using the coloc2 algorithm in FIJI image analysis software. This algorithm was used to calculate the Pearson coefficient. The results from the coloc2-derived Pearson coefficient were graphed using GraphPad Prism 10.0 (GraphPad Software, Inc.) with statistical significance established at $p < 0.05$.

Fluorescence Lifetime Measurement (FLIM)
CHO cells (60,000 cells) seeded on a 33 mm glass-bottom dish (MatTek) were transfected using Lipofectamine 3000 (Invitrogen) with mNeonGreen-mCherry dimer, co-transfected with FMRP-mCherry and shank2-mNeonGreen, or FMRP-mCherry and shank2-mNeonGreen mutants (R958S and A1731S). 36 hours following transfection cells were imaged in a temperature-controlled incubator 37 °C supplemented with 5% $CO_2$. mNeonGreen fluorescent lifetime ($\tau$; tau; expressed in ns) was determined by stimulation with a 440 nm pulse laser and imaged on Nikon A1R confocal Microscope using a 40x / 1.25 objective. PicoQuant Software SymPhotime (Germany) captured 3x digital zooms of fluorescent positive CHO cells and was used to calculate $\tau$ values. mNeonGreen donor alone was used to determine basal (control) lifetimes. mNeonGreen-mCherry dimer was used as a Förster resonance energy transfer (FRET) positive control. FMRP-mCherry was used as an acceptor to quantitate FRET seen as a reduction in shank2-mNeonGreen and shank2-mNeonGreen mutants fluorescence timescales, respectively. Lifetime decay slopes ($\tau$) were derived from 3 to 12 ns following the laser pulse and $\Delta\tau$ were determined by basal $\tau$ minus measured $\tau$ values. Lifetime values were statistically compared using 1-way ANOVA.

Western Blots of FMRP, pFMRP, PP2A subunit C, and CaMKII
Protein was extracted from cells grown in 6-well plates by washing the cells 2 times with ice-cold PBS, followed by immersion in RIPA buffer (150 mM sodium chloride, 1.0 % NP-40 or Triton X-100, 0.5 % sodium deoxycholate, 0.1 % SDS, 50 mM Tris, pH 8.0) with a complete Mini protease inhibitor. The cells in RIPA buffer were kept on ice for 10 minutes and vortexed every 2–3 minutes. After 10 minutes, the samples were sonicated on a Fisher Scientific sonic dismembrator F-60 and stored at −80 °C. Proteins were quantified using the BioRad DC Assay kit. At least 20 µg of protein was loaded per sample per lane of a BOLT 4–12 % gradient gel loaded into a BOLT mini gel tank. Proteins were transferred overnight onto nitrocellulose membrane at 60 V. The membranes were blocked in either 5 % BSA in Tris-buffered saline with 0.05 % Tween 20 (TBS-T) or 5 % milk in phosphate-buffered saline with 0.05% Tween 20 (PBS-T) for 1 hour at room temperature (22 °C). Membranes were incubated with primary antibodies (1:1,000 to 1:5,000 dilution depending on antibody) overnight at 4 °C. Primary antibodies include Shank2 (Cell Signaling Technologies), FMRP (Abcam), pFMRP S499

(Abcam), PP2A subunit C (Sigma), and CaMKIIα (Abcam), β-actin (Sigma), and GAPDH (Sigma). The following day, the membranes were washed 3 times for 5 minutes and incubated with HRP-conjugated secondary antibodies (1:5,000) for 1 hour at room temperature. The membranes were then washed 3 times for 5 minutes. To visualize protein, either SuperSignal West Pico Chemiluminescent Substrate (used for loading controls) or SuperSignal West Femto Maximum Sensitivity Substrate (used for all other proteins) was applied to the membrane as evenly as possible. Shortly thereafter the membranes were exposed and imaged on a Biorad Gel Doc XR system. The resulting images were quantified using BioRad ImageLab 5.1.4 software.

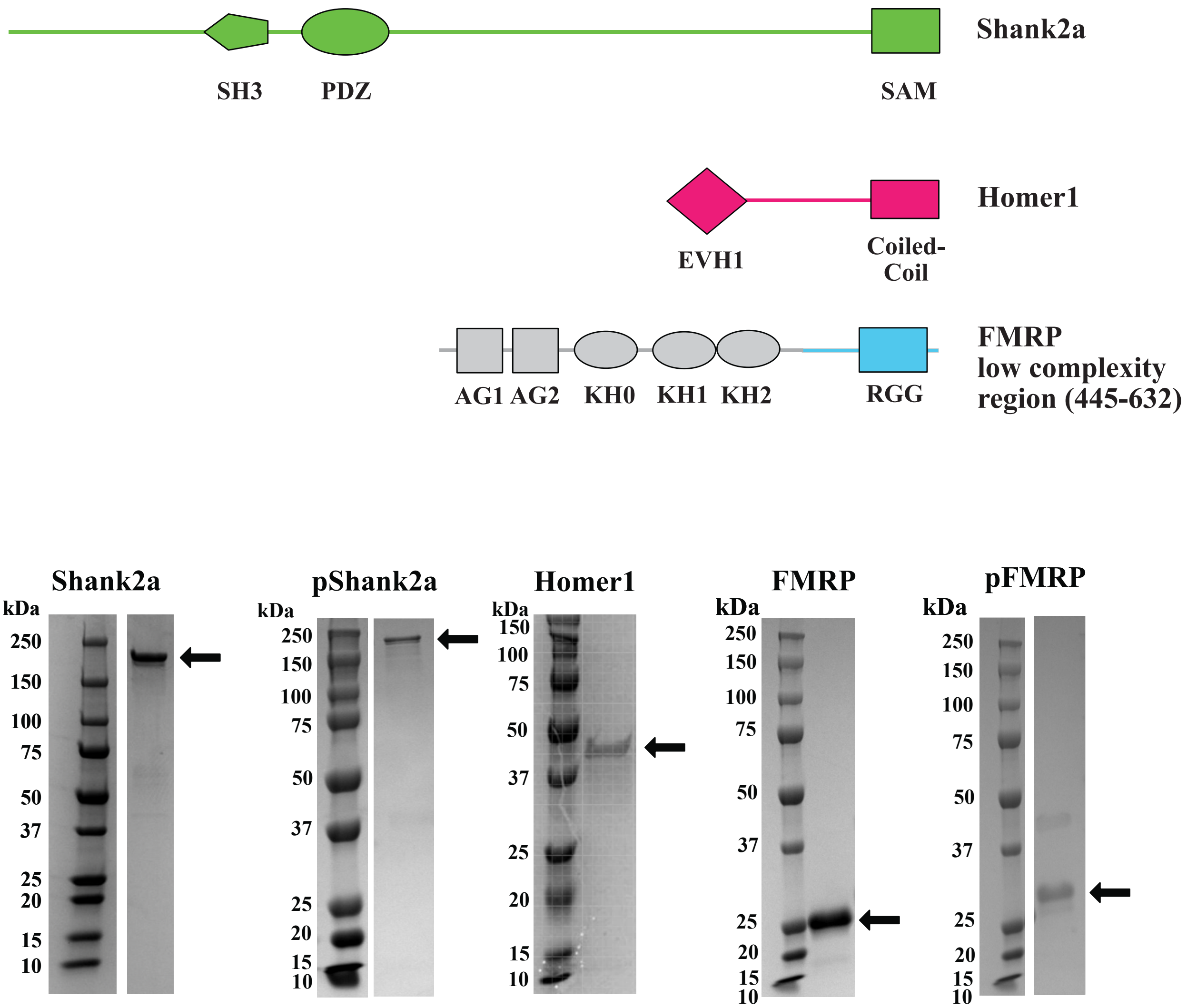


**Fig. S1. Recombinant proteins used in biochemical assays.** (top) Schematics of shank2, homer1, and FMRP purified and used in biochemical assays. SH3 = Src homology 3 domain. PDZ = Postsynaptic density protein, disc large tumor suppressor, zonula occludens-1 protein domain. SAM = Sterile alpha motif. EVH1 = enabled/vasodilator-stimulated phosphoprotein homology 1 domain. Colored (green, magenta, cyan) portions of protein indicate expressed and purified regions. (bottom) Coomassie-stained SDS-PAGE gels of purified shank2, homer1, and FMRP, and phosphorylated shank2 and FMRP.

| | | | | |
|---|---|---|---|---|
| MKSLLNAFTK | KEVPFREAPA | YSNRRRRPPN | TLAAPRVLLR | SNSDNNLNAS |
| APDWAVCSTA | TSHRSLSPQL | LQQMPSKPEG | AAKTIGSYVP | GPRSRSPSLN |
| RLGGAGEDGK | RPQPLWHVGS | PFALGANKDS | LSAFEYPGPK | RKLYSAVPGR |
| LFVAVKPYQP | QVDGEIPLHR | GDRVKVLSIG | EGGFWEGSAR | GHIGWFPAEC |
| VEEVQCKPRD | SQAETRADRS | KKLFRHYTVG | SYDSFDTSSD | CIIEEKTVVL |
| QKKDNEGFGF | VLRGAKADTP | IEEFTPTPAF | PALQYLESVD | EGGVAWQAGL |
| RTGDFLIEVN | NENVVKVGHR | QVVNMIRQGG | NHLVLKVVTV | TRNLDPDDTA |
| RKKAPPPPKR | APTTALTLRS | KSMTSELEEL | VDKASVRKKK | DKPEEIVPAS |
| KPSRAAENMA | VEPRVATIKQ | RPSSRCFPAG | SDMNSVYERQ | GIAVMTPTVP |
| GSPKAPFLGI | PRGTMRRQKS | SRIFLSGI | TEEERQFLAP | PMLKFTRSLS |
| MPDTSEDIPP | PPQSVPPSPP | PPSPTTYNCP | KSPTPRVYGT | IKPAFNQNSA |
| AKVSPATRSD | TVATMMREKG | MYFRRELDRY | SLDSEDLYSR | NAGPQANFRN |
| KRGQMPENPY | SEVGKIASKA | VYVPAKPARR | KGMLVKQSNV | EDSPEKTCSI |
| PIPTIIVKEP | STSSSGKSSQ | GSSMEIDPQA | PEPPSQLRPD | ESLTVSSPFA |
| AAIAGAVRDR | EKRLEARRNS | PAFLSTDLGD | EDVGLGPPAP | RTRPSMFPEE |
| GDFADEDSAE | QLSSPMPSAT | PREPENHFVG | GAEASAPGEA | GRPLNSTSKA |
| QGPESSPAVP | SASSGTAGPG | NYVHPLTGRL | LDPSSPLAL A | LSARDRAMKE |
| SQQGPKGEAP | KADLNKPLYI | DTKMRPSLDA | GFPTVTRQNT | RGPLRRQETE |
| NKYETDLGRD | RKGDDKKNML | IDIMDTSQQK | SAGLLMVHTV | DATKLDNALQ |
| EEDEKAEVEM | KPDSSPSEVP | EGVSETEGAL | QISAAPEPTT | VPGRTIVAVG |
| SMEEAVILPF | RIPPPPLASV | DLDEDFIFTE | PLPPPLEFAN | SFDIPDDRAA |
| SVPALSDLVK | QKKSDTPQSP | SLNSSQPTNS | ADSKKPASLS | NCLPASFLPP |
| PESFDAVADS | GIEEVDSRSS | SDHHLETTST | ISTVSSISTL | SSEGGENVDT |
| CTVYADGQAF | MVDKPPVPPK | PKMKPIIHKS | NALYQDALVE | EDVDSFVIPP |
| PAPPPPPGSA | QPGMAKVLQP | RTSKLWGDVT | EIKSPILSGP | KANVISELNS |
| ILQQMNREKL | AKPGEGLDSP | MGAKSASLAP | RSPEIMSTIS | GTRSTTVTFT |
| VRPGTSQPIT | LQSRPPDYES | RTSGTRRAPS | PVVSPTEMNK | ETLPAPLSAA |
| TASPSPALSD | VFSLPSQPPS | GDLFGLNPAG | RSRSPSPSIL | QQPISNKPFT |
| TKPVHLWTKP | DVADWLESLN | LGEHKEAFMD | NEIDGSHLPN | LQKEDLIDLG |
| VTRVGHRMNI | ERALKQLLDR | | | |

Src homology 3 domain

Postsynaptic density protein, disc large tumor suppressor, zonula occludens-1 protein domain

Sterile Alpha Motif

▽ Mass-Spec detected phosphorylation site

**Fig. S2. Mass-spectometry analysis of phosphorylated shank2.** Mitogen-activated protein kinase phosphorylates shank2 on 10 sites, 7 of which were in the intrinsically disordered region. It is important to note that this analysis detected sites across proteins in solution and does not indicate that each shank2 molecule was phosphorylated on 10 sites. There is likely varying degrees of phosphorylation on individual shank2 proteins.

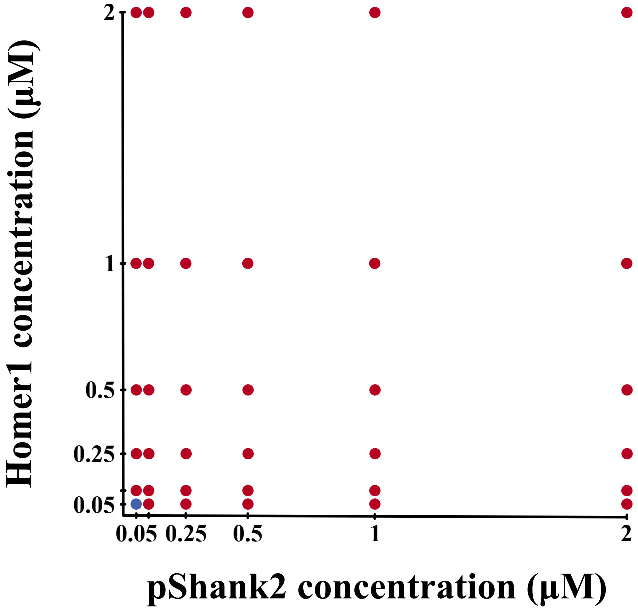


**Fig. S3. Phase diagram of phosphorylated shank2 and homer1.** Phase diagram of phosphorylated shank2 and homer1 as concentrations of each increased from 0.05 µM to 5 µM. Blue dots indicate no observable phase separation; red dots indicate phase separation. N = 3 experiments with 5 fields of view each analyzed.

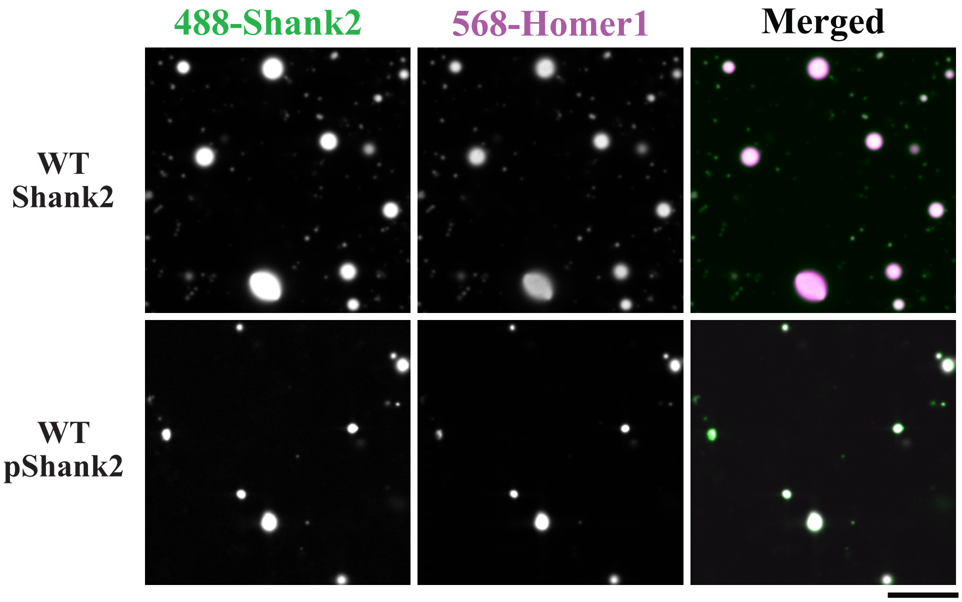


**Fig. S4. Shank2 and homer1 condensates form in rabbit reticulocyte lysates.** Condensates composed of 0.5 µM shank2 (green) and 0.5 µM homer1 (magenta) that are formed in rabbit reticulocyte lysate are similar to those observed in solution-based assays shown in Fig. 1. Scale bar = 10 µm.

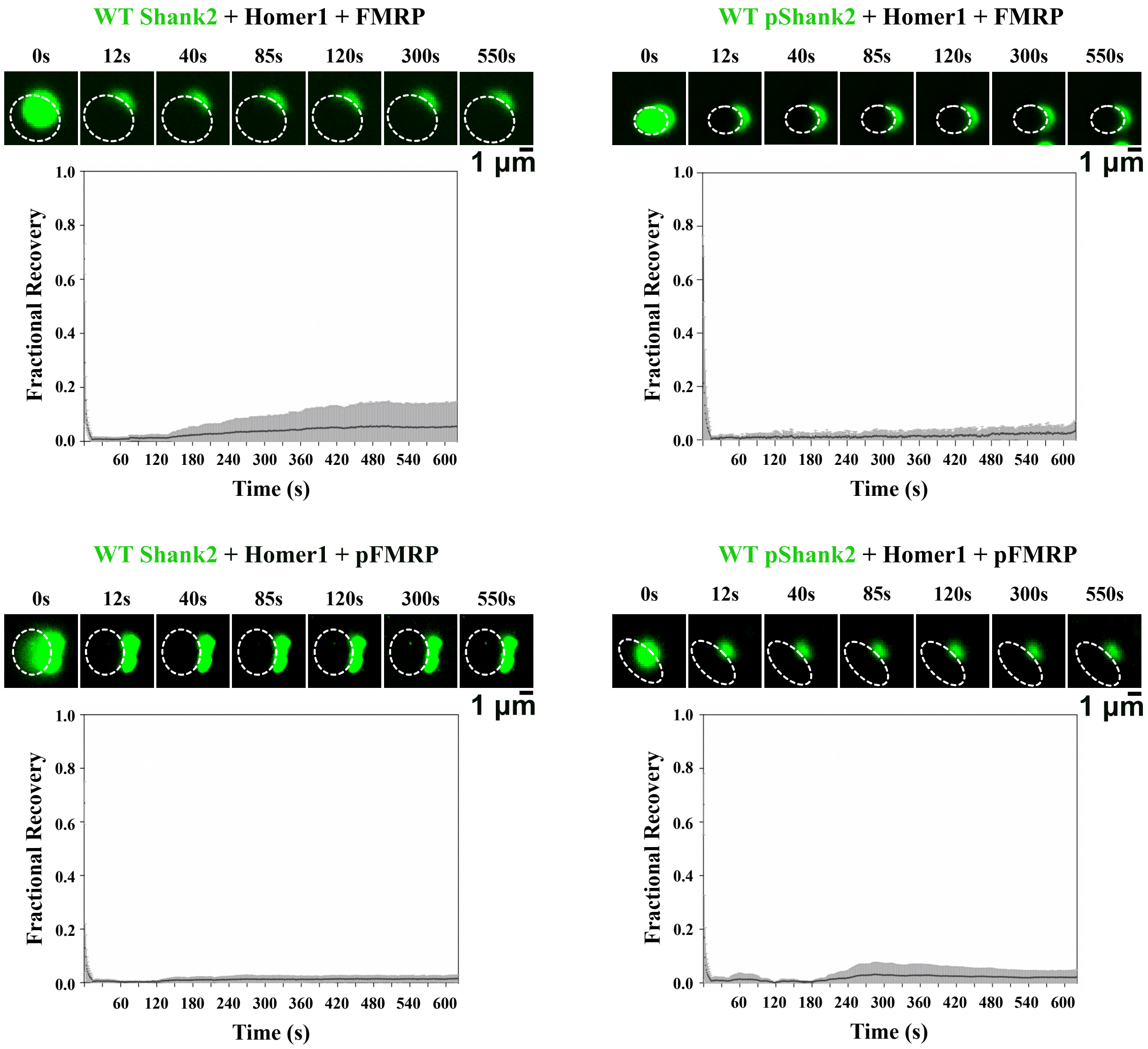


**Fig. S5. Shank2 dynamics in shank2, homer1, and FMRP-enriched condensates are arrested.** Fluorescence recovery after photobleaching (FRAP) of AF488-shank2 (green) and homer1 condensate. Time 0 s was captured before photobleaching, 12 s was captured immediately after the photobleaching pulse. Dashed white box illustrates region of photobleach. Scale bar = 1 μm. Plot on bottom shows the time course of recovered for AF488-shank2 in condensates formed with shank2 and homer1. Shown are the mean (black line) ± s.d. (gray region above and below black line). N = 3.

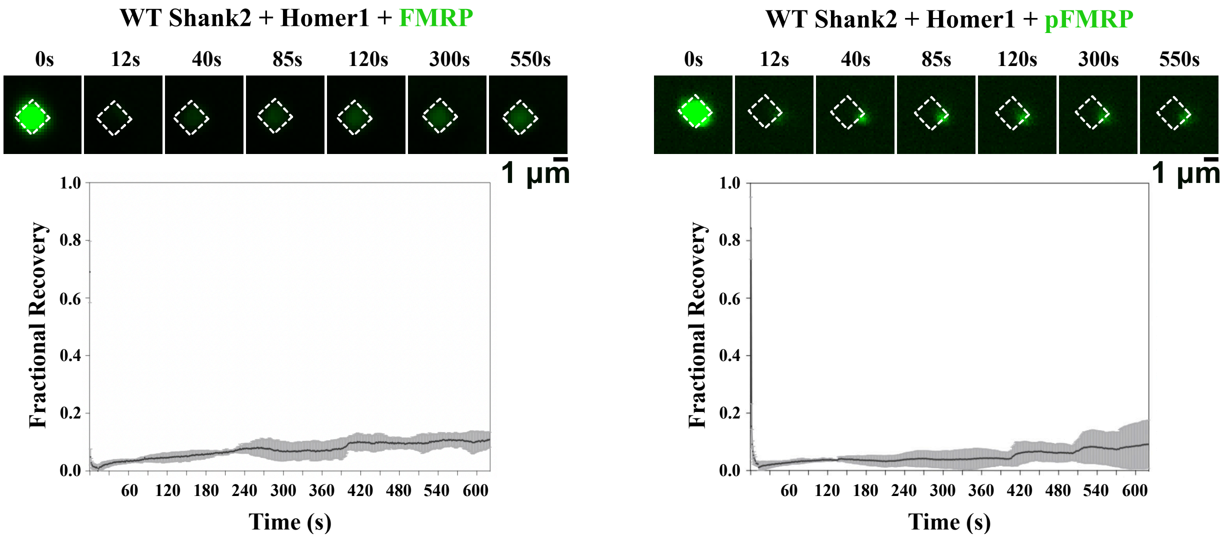


**Fig. S6. FMRP dynamics in shank2, homer1, and FMRP-enriched condensates are arrested.** Fluorescence recovery after photobleaching (FRAP) of AF488-FMRP (green) in shank2, homer1, and FMRP-enriched condensates. Time 0 s was captured before photobleaching, 12 s was captured immediately after the photobleaching pulse. Dashed white box illustrates region of photobleach. Scale bar = 1 µm. Plot on bottom shows the time course of recovered for AF488-shank2 in condensates formed with shank2 and homer1. Shown are the mean (black line) ± s.d. (gray region above and below black line). N = 3.

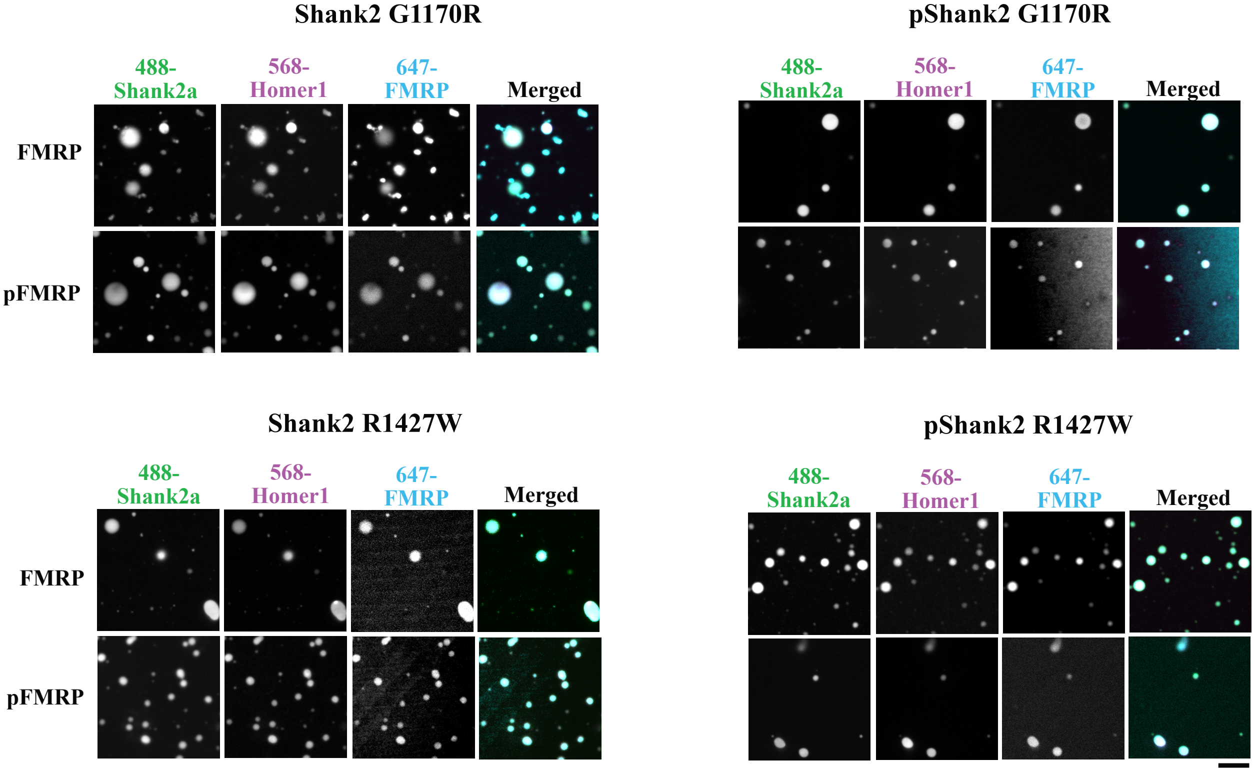


**Fig. S7. Neither G1170R nor R1427W shank2 mutations affect FMRP enrichment in shank2 and homer1 condensates.** AF647-FMRP (cyan) or AF647-phospho-FMRP (pFMRP, cyan) are enriched in AF488-shank2 (green) / AF568-homer1 (magenta) condensates or AF488-phospho-shank2 (pshank2, green) / AF468-homer1 (magenta) condensates. Scale bar = 10 µm. N = 3 experiments with 5 fields of view each analyzed.

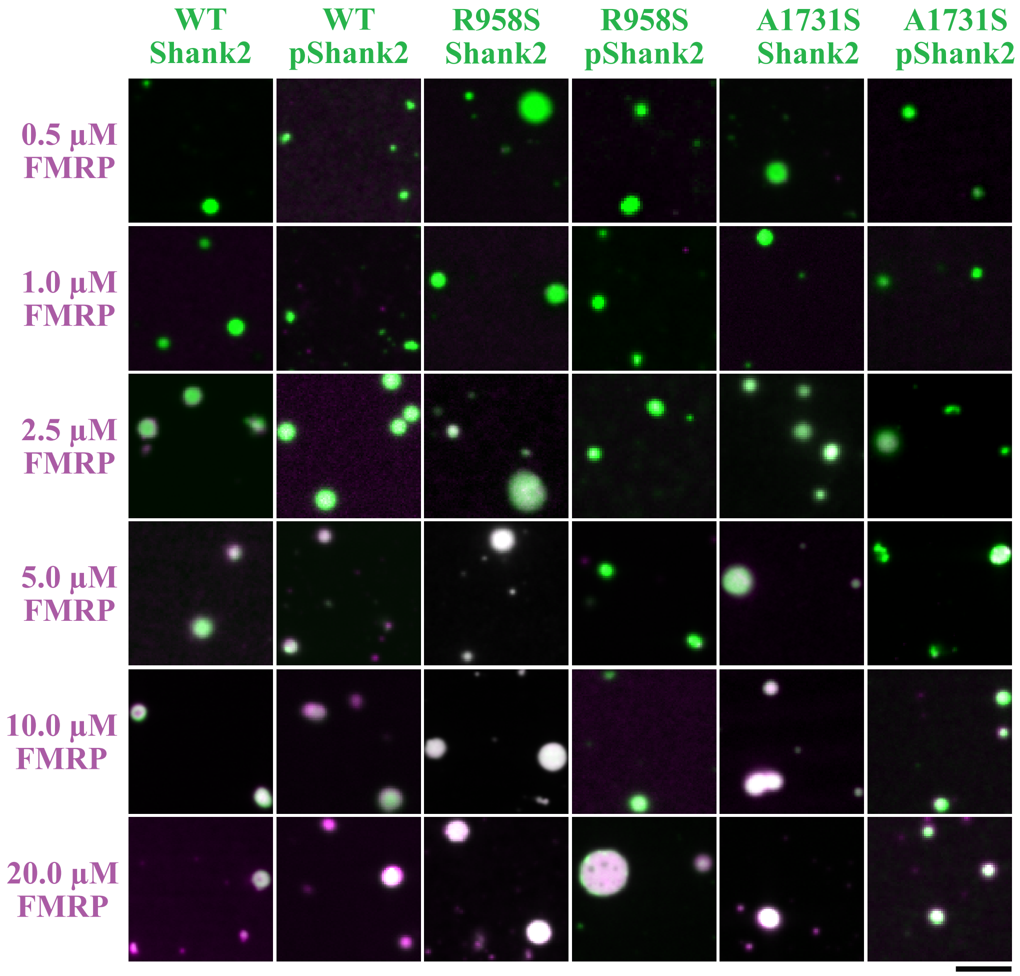


**Fig. S8. FMRP enrichment in R958S and A1731S shank2 and homer1 condensates.** FMRP (magenta) concentration was increased from 0.5 µM to 20 µM in the presence of (p)shank2 (green, 0.5 µM) and homer1 (0.5 µM) condensates. As the concentration of FMRP increases, enrichment in shank2 and homer1 condensates increases, as indicated by increasing white in condensates in the merged images. Quantification of images is shown in Fig. 3C. Scale bar = 5 µm. N = 3 experiments with 5 fields of view each analyzed.

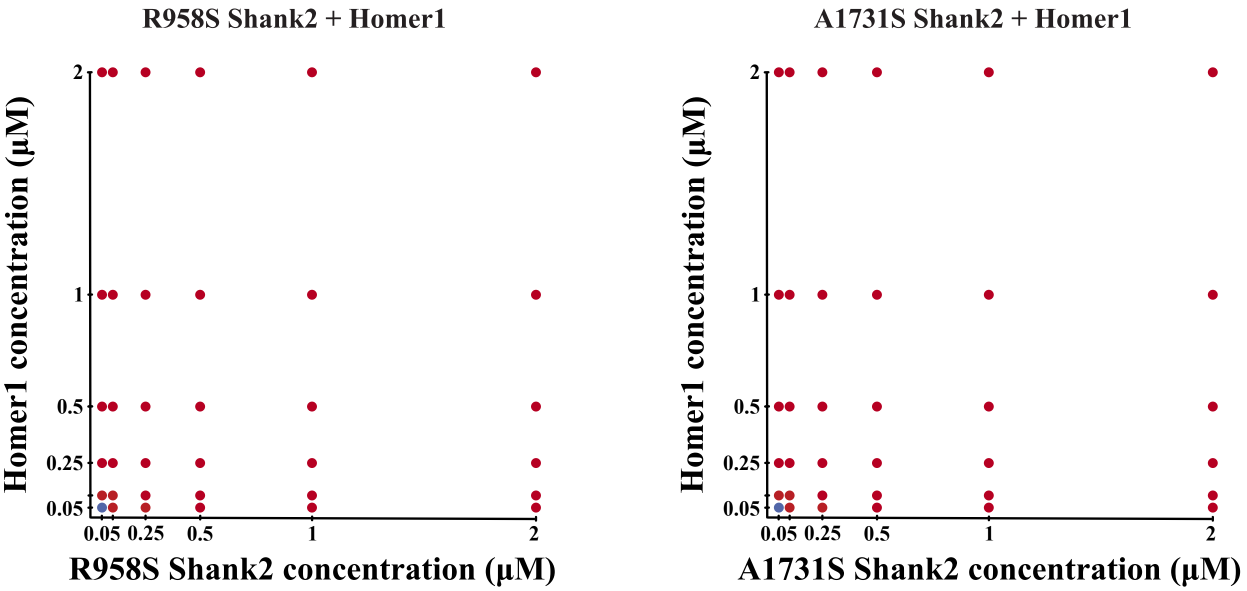


**Fig. S9. Neither R958S nor A1731S shank2 mutations affect the critical concentrations for phase separation of mutant shank2 and homer1.** Phase diagrams of R958S or A1731S shank2 and homer1 as concentrations of each increased from 0.05 µM to 5 µM. Blue dots indicate no observable phase separation; red dots indicate phase separation. N = 3 experiments with 5 fields of view each analyzed.

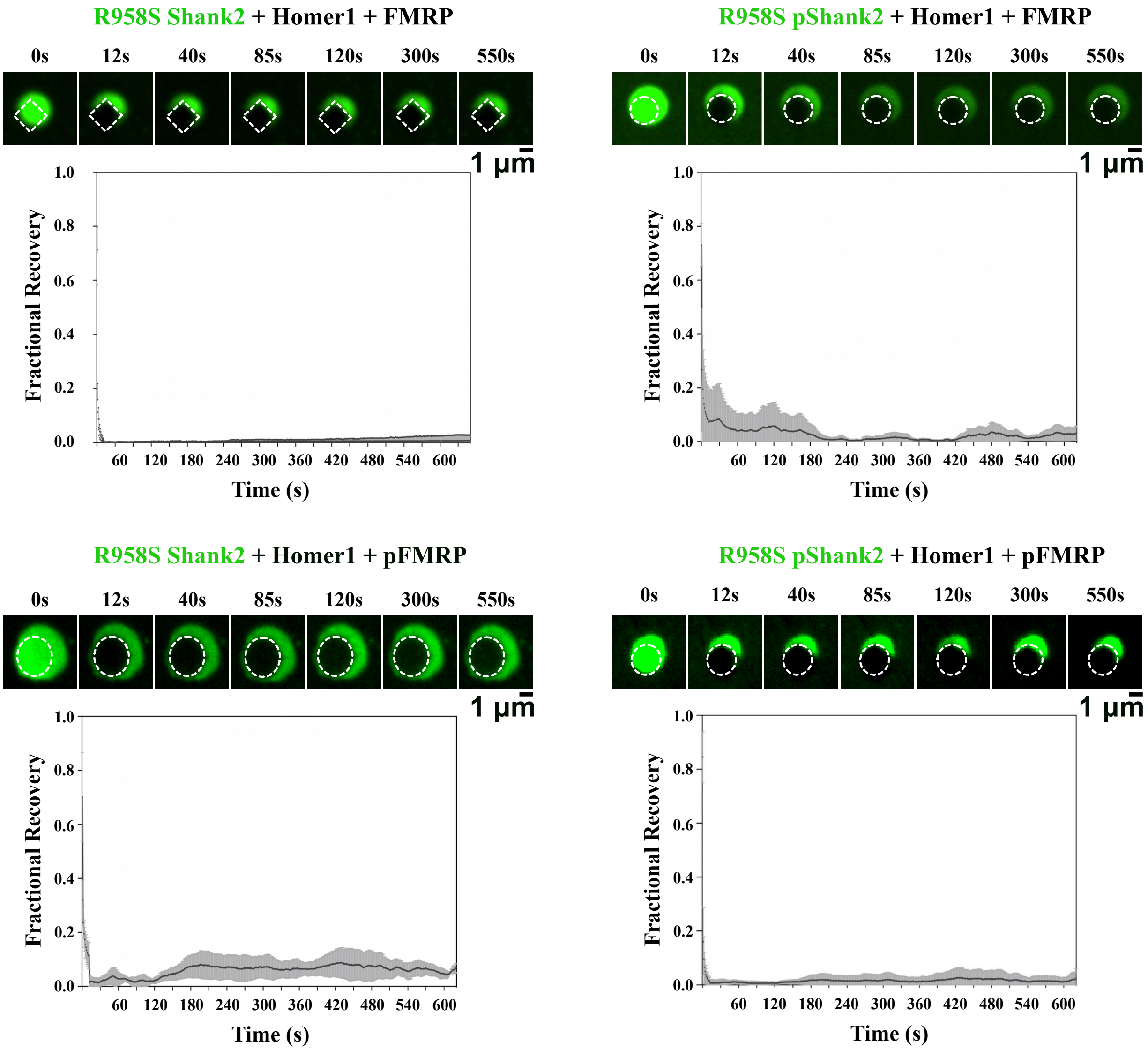


**Fig. S10. R958S shank2 mutations alter arrested shank2 dynamics in shank2, homer1, and FMRP-enriched condensates.** Fluorescence recovery after photobleaching (FRAP) of AF488-R958S shank2 or AF488-R958S pshank2 (green) in shank2, homer1, and (p)FMRP-enriched condensates. Time 0 s was captured before photobleaching, 12 s was captured immediately after the photobleaching pulse. Dashed white box illustrates region of photobleach. Scale bar = 1 µm. Plot on bottom shows the time course of recovered for AF488-shank2 in condensates formed with shank2 and homer1 and enriched with FMRP or pFMRP. Shown are the mean (black line) ± s.d. (gray region above and below black line). N = 3.

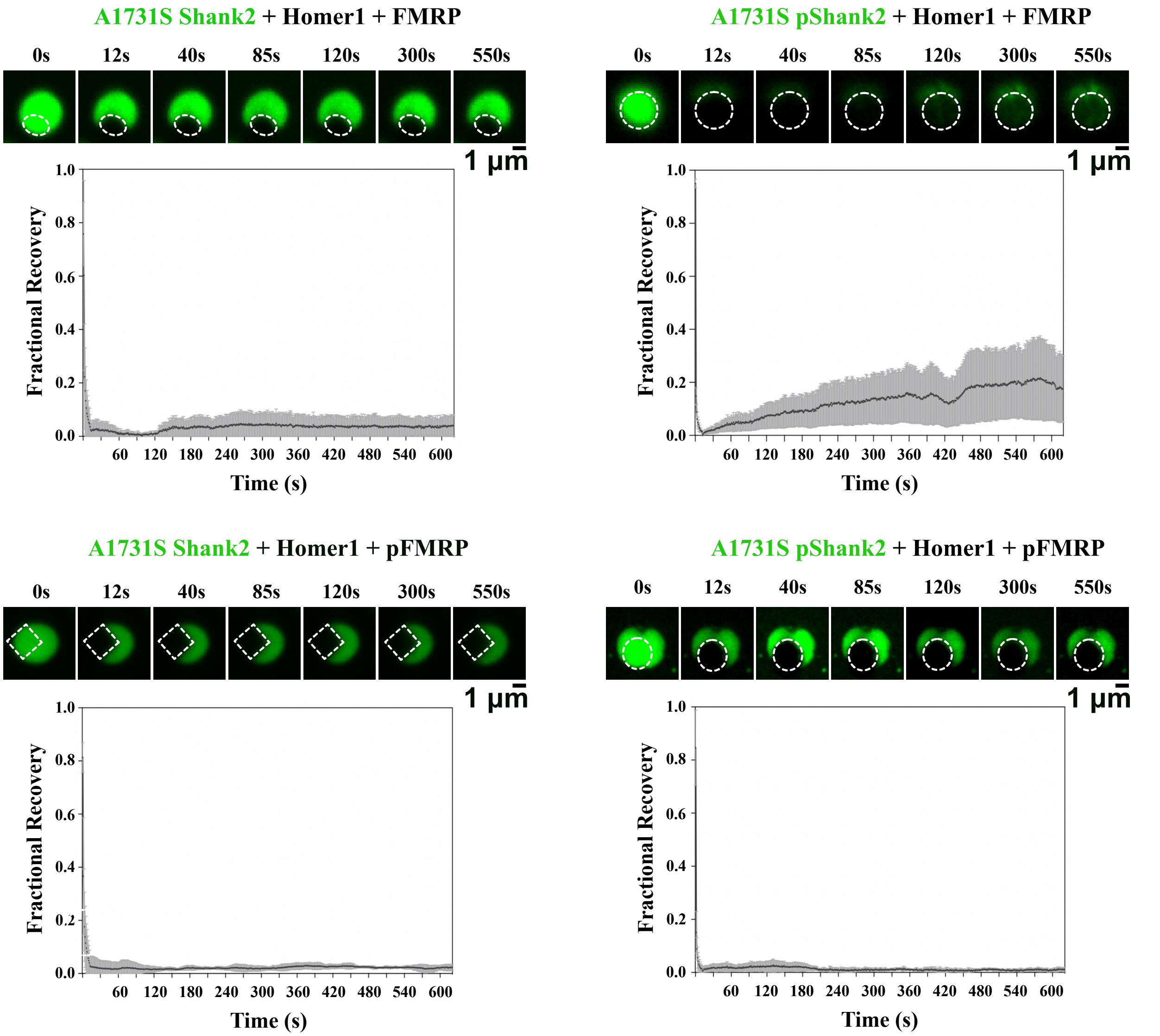


**Fig. S11. A1731S pshank2 dynamics are slightly increased at the edges of A1731S pshank2, homer1, and FMRP-enriched condensates.** Fluorescence recovery after photobleaching (FRAP) of AF488-A1731S shank2 or AF488-A1731S pshank2 (green) in shank2, homer1, and (p)FMRP-enriched condensates. Time 0 s was captured before photobleaching, 12 s was captured immediately after the photobleaching pulse. Dashed white box illustrates region of photobleach. Scale bar = 1 µm. Plot on bottom shows the time course of recovered for AF488-shank2 in condensates formed with shank2 and homer1 and enriched with FMRP or pFMRP. Shown are the mean (black line) ± s.d. (gray region above and below black line). N = 3.

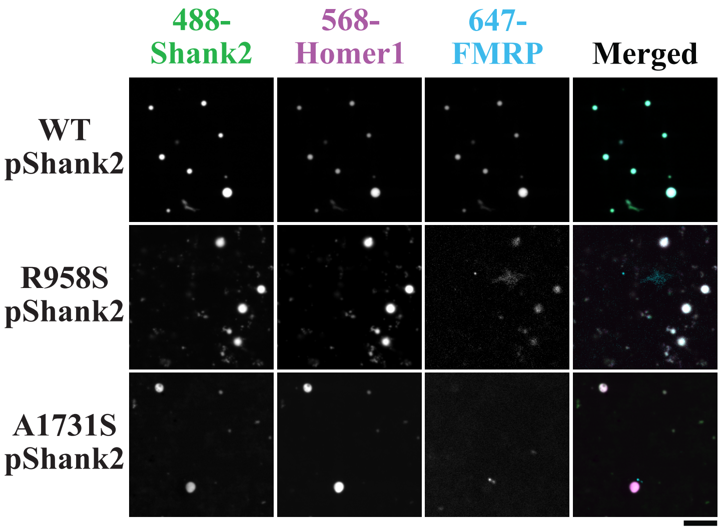


**Fig. S12. Enrichment of FMRP is diminished in R958S and A1731S shank2 and homer1 condensates compared with wild type shank2 and homer1 condensates in rabbit reticulocyte lysate.** These results are similar to what was observed in solution-based assays shown in Fig. 3A. Scale bar = 10 µm. N = 3 experiments with 5 fields of view each analyzed.

**Table S1. Data set for Shear Rate measurements shown in Fig. 3E.**

| | WT pShank2 | | | R958S pShank2 | | | A1731S pShank2 | |
|---|---|---|---|---|---|---|---|---|
| **Shear Rate γ ($s^{-1}$)** | **Average Viscosity (Pa.s)** | **StDev Viscosity (Pa.s)** | | **Average Viscosity (Pa.s)** | **StDev Viscosity (Pa.s)** | | **Average Viscosity (Pa.s)** | **StDev Viscosity (Pa.s)** |
| 10 | 0.0228 | 0.0051 | | 0.0876 | 0.0440 | | 0.0680 | 0.0457 |
| 16 | 0.0080 | 0.0029 | | 0.0541 | 0.0280 | | 0.0526 | 0.0276 |
| 25 | 0.0051 | 0.0025 | | 0.0312 | 0.0145 | | 0.0393 | 0.0171 |
| 40 | 0.0044 | 0.0022 | | 0.0202 | 0.0079 | | 0.0261 | 0.0098 |
| 63 | 0.0038 | 0.0017 | | 0.0141 | 0.0041 | | 0.0166 | 0.0048 |
| 100 | 0.0035 | 0.0010 | | 0.0102 | 0.0020 | | 0.0110 | 0.0024 |
| 158 | 0.0032 | 0.0004 | | 0.0073 | 0.0009 | | 0.0072 | 0.0006 |
| 251 | 0.0026 | 0.0003 | | 0.0058 | 0.0011 | | 0.0054 | 0.0005 |
| 398 | 0.0022 | 0.0003 | | 0.0044 | 0.0008 | | 0.0042 | 0.0008 |
| 631 | 0.0019 | 0.0003 | | 0.0034 | 0.0006 | | 0.0033 | 0.0009 |
| 1000 | 0.0016 | 0.0004 | | 0.0027 | 0.0004 | | 0.0027 | 0.0008 |

**Table S2. Data set for Storage Modulus measurements shown in Fig. 3E.**

| | WT pShank2 | | | R958S pShank2 | | | A1731S pShank2 | |
|---|---|---|---|---|---|---|---|---|
| **Angular Frequency ω (rad / s)** | **Average Storage Modulus G' (Pa)** | **StDev Storage Modulus G' (Pa)** | | **Average Storage Modulus G' (Pa)** | **StDev Storage Modulus G' (Pa)** | | **Average Storage Modulus G' (Pa)** | **StDev Storage Modulus G' (Pa)** |
| 1 | 0.1081 | 0.0651 | | 0.8670 | 0.1508 | | 0.4221 | 0.2287 |
| 1.58 | 0.1226 | 0.0708 | | 0.9684 | 0.2977 | | 0.5447 | 0.3492 |
| 2.51 | 0.1457 | 0.0819 | | 1.1104 | 0.4771 | | 0.6488 | 0.4408 |
| 3.98 | 0.2000 | 0.0994 | | 1.2833 | 0.6788 | | 0.7858 | 0.5448 |
| 6.31 | 0.2849 | 0.1236 | | 1.5220 | 0.9014 | | 0.9459 | 0.6531 |
| 10 | 0.4097 | 0.1406 | | 1.8473 | 1.1571 | | 1.1477 | 0.7807 |
| 15.85 | 0.5585 | 0.1604 | | 2.2492 | 1.4745 | | 1.3932 | 0.9399 |
| 25.11 | 0.7551 | 0.1955 | | 2.7462 | 1.8321 | | 1.6934 | 1.0991 |
| 39.82 | 1.0682 | 0.3123 | | 3.2717 | 2.0579 | | 2.1118 | 1.1530 |
| 63.10 | 1.8859 | 0.7707 | | 3.7738 | 1.8099 | | 2.9197 | 0.9274 |
| 100 | 3.7174 | 1.8470 | | 4.4613 | 2.3358 | | 5.0889 | 1.1201 |

**Table S3. Data set for Loss Modulus measurements shown in Fig. 3E.**

| | WT pShank2 | | | R958S pShank2 | | | A1731S pShank2 | |
|---|---|---|---|---|---|---|---|---|
| Angular Frequency ω (rad / s) | Average Storage Modulus G' (Pa) | StDev Storage Modulus G' (Pa) | | Average Storage Modulus G' (Pa) | StDev Storage Modulus G' (Pa) | | Average Storage Modulus G' (Pa) | StDev Storage Modulus G' (Pa) |
| 1 | 0.0552 | 0.0363 | | 0.1716 | 0.0258 | | 0.1840 | 0.0897 |
| 1.58 | 0.0543 | 0.0346 | | 0.1753 | 0.0312 | | 0.1937 | 0.0990 |
| 2.51 | 0.0544 | 0.0326 | | 0.1871 | 0.0429 | | 0.2029 | 0.1034 |
| 3.98 | 0.0566 | 0.0307 | | 0.2006 | 0.0503 | | 0.2162 | 0.1106 |
| 6.31 | 0.0598 | 0.0274 | | 0.2191 | 0.0575 | | 0.2351 | 0.1267 |
| 10 | 0.0665 | 0.0235 | | 0.2408 | 0.0654 | | 0.2598 | 0.1512 |
| 15.85 | 0.0737 | 0.0172 | | 0.2676 | 0.0754 | | 0.2894 | 0.1854 |
| 25.11 | 0.0728 | 0.0151 | | 0.3103 | 0.1088 | | 0.3224 | 0.2391 |
| 39.82 | 0.0407 | 0.0454 | | 0.3461 | 0.1643 | | 0.3333 | 0.3123 |
| 63.10 | | | | 0.2902 | 0.2204 | | 0.2433 | 0.3840 |
| 100 | | | | | | | | |

**Table S4. Sequences of proteins used in biochemical assays in this study.**

| **Protein** | **Sequence** | **Notes** |
|---|---|---|
| Shank2a | **MGSSHHHHHHGSGLVPRGSASMSDSEVNQE**<br>**AKPEVKPEVKPETHINLKVSDGSSEIFFKIKK**<br>**TTPLRRLMEAFAKRQGKEMDSLRFLYDGIRI**<br>**QADQTPEDLDMEDNDIIEAHREQIGG**MKSLL<br>NAFTKKEVPFREAPAYSNRRRRPPNTLAAPRVL<br>LRSNSDNNLNASAPDWAVCSTATSHRSLSPQLL<br>QQMPSKPEGAAKTIGSYVPGPRSRSPSLNRLGG<br>AGEDGKRPQPLWHVGSPFALGANKDSLSAFEY<br>PGPKRKLYSAVPGRLFVAVKPYQPQVDGEIPLH<br>RGDRVKVLSIGEGGFWEGSARGHIGWFPAECV<br>EEVQCKPRDSQAETRADRSKKLFRHYTVGSYD<br>SFDTSSDCIIEEKTVVLQKKDNEGFGFVLRGAK<br>ADTPIEEFTPTPAFPALQYLESVDEGGVAWQAG<br>LRTGDFLIEVNNENVVKVGHRQVVNMIRQGGN<br>HLVLKVVTVTRNLDPDDTARKKAPPPPKRAPTT<br>ALTLRSKSMTSELEELVDKASVRKKKDKPEEIV<br>PASKPSRAAENMAVEPRVATIKQRPSSRCFPAG<br>SDMNSVYERQGIAVMTPTVPGSPKAPFLGIPRG<br>TMRRQKSIDSRIFLSGITEEERQFLAPPMLKFTRS<br>LSMPDTSEDIPPPPQSVPPSPPPPSPTTYNCPKSPT<br>PRVYGTIKPAFNQNSAAKVSPATRSDTVATMM<br>REKGMYFRRELDRYSLDSEDLYSRNAGPQANF<br>RNKRGQMPENPYSEVGKIASKAVYVPAKPARR<br>KGMLVKQSNVEDSPEKTCSIPIPTIIVKEPSTSSS<br>GKSSQGSSMEIDPQAPEPPSQLRPDESLTVSSPFA<br>AAIAGAVRDREKRLEARRNSPAFLSTDLGDEDV<br>GLGPPAPRTRPSMFPEEGDFADEDSAEQLSSPMP<br>SATPREPENHFVGGAEASAPGEAGRPLNSTSKA<br>QGPESSPAVPSASSGTAGPGNYVHPLTGRLLDPS<br>SPLALALSARDRAMKESQQGPKGEAPKADLNK<br>PLYIDTKMRPSLDAGFPTVTRQNTRGPLRRQET<br>ENKYETDLGRDRKGDDKKNMLIDIMDTSQQKS<br>AGLLMVHTVDATKLDNALQEEDEKAEVEMKP<br>DSSPSEVPEGVSETEGALQISAAPEPTTVPGRTIV<br>AVGSMEEAVILPFRIPPPPLASVDLDEDFIFTEPL<br>PPPLEFANSFDIPDDRAASVPALSDLVKQKKSDT<br>PQSPSLNSSQPTNSADSKKPASLSNCLPASFLPPP<br>ESFDAVADSGIEEVDSRSSSDHHLETTSTISTVSS<br>ISTLSSEGGENVDTCTVYADGQAFMVDKPPVPP<br>KPKMKPIIHKSNALYQDALVEEDVDSFVIPPPAP<br>PPPPGSAQPGMAKVLQPRTSKLWGDVTEIKSPIL<br>SGPKANVISELNSILQQMNREKLAKPGEGLDSP<br>MGAKSASLAPRSPEIMSTISGTRSTTVTFTVRPG<br>TSQPITLQSRPPDYESRTSGTRRAPSPVVSPTEM | Human shank2a with N-terminal SUMO tag. Shank2a isoform consists of residues 391 – 1849 of full length shank2 with an additional 12-residue MKSLLNAFTKKE sequence at the N-terminus. SUMO sequence is **blue**.<br><br>Site-directed mutagenesis was used to generate R958S, G1170R, R1427W, and A1731S mutations. |

| | | |
|---|---|---|
| | NKETLPAPLSAATASPSPALSDVFSLPSQPPSGD<br>LFGLNPAGRSRSPSPSILQQPISNKPFTTKPVHLW<br>TKPDVADWLESLNLGEHKEAFMDNEIDGSHLP<br>NLQKEDLIDLGVTRVGHRMNIERALKQLLDR | |
| Homer1 | MGEQPIFSTRAHVFQIDPNTKKNWVPTSKHAVT<br>VSYFYDSTRNVYRIISLDGSKAIINSTITPNMTFT<br>KTSQKFGQWADSRANTVYGLGFSSEHHLSKFA<br>EKFQEFKEAARLAKEKSQEKMELTSTPSQESAG<br>GDLQSPLTPESINGTDDERTPDVTQNSEPRAEPT<br>QNALPFSHSSAISKHWEAELATLKGNNAKLTAA<br>LLESTANVKQWKQQLAAYQEEAERLHKRVTEL<br>ECVSSQANAVHTHKTELNQTIQELEETLKLKEE<br>EIERLKQEIDNARELQEQRDSLTQKLQEVEIRNK<br>DLEGQLSDLEQRLEKSQNEQEAFRNNLKTLLEI<br>LDGKIFELTELRDNLAKLLECS | Full-length human homer1 sequence. Protein is expressed with an N-terminal His$_6$-SUMO tag that was cleaved using ULP1 protease. |
| FMRP C-terminal IDR | GASSRPPPNRTDKEKSYVTDDGQGMGRGSRPYR<br>NRGHGRRGPGYTSGTNSEASNASETESDHRDELS<br>DWSLAPTEEERESFLRRGDGRRRGGGGRGQGGR<br>GRGGGFKGNDDHSRTDNRPRNPREAKGRTTDGS<br>LQIRVDCNNERSVHTKTLQNTSSEGSRLRTGKDR<br>NQKKEKPDSVDGQQPLVNGVP | Human FMRP residues 445-632. Expressed as a His$_6$-SUMO-FMRP$_{445\text{-}632}$ fusion. SUMO was cleaved using ULP1 protease. |

**Author Contributions**

L.A.L and J.A.D. conceived of the project and designed the experiments with input for neuronal experiments from W.W., F.M., and J.E. L.A.L, S.C., and X.L. prepared biochemical reagents. L.A.L. and W.W. differentiated and cultured human neurons from NPCs. L.A.L. performed all microscopy-based biochemical assays, *in vitro* RNA translation assays, and neuronal transfections and imaging. S.C. and X.L. performed rheometry experiments. J.A. performed CHO cell culture and FLIM assays. F.M. differentiated and cultured wild type, knock out, and R841X shank2 neurons and performed FMRP, pFMRP, and CaMKII detection by western blot and CaMKII RNA counts by RNA Seq experiments. L.A.L. and J.A.D. wrote the manuscript with input from J.E., J.A., S.C. W.W., F.M., and X.L.
